# Evolutionary systems biology reveals patterns of rice adaptation to drought-prone agro-ecosystems

**DOI:** 10.1101/2021.05.26.445872

**Authors:** Simon C. Groen, Zoé Joly-Lopez, Adrian E. Platts, Mignon Natividad, Zoë Fresquez, William M. Mauck, Marinell R. Quintana, Carlo Leo U. Cabral, Rolando O. Torres, Rahul Satija, Michael D. Purugganan, Amelia Henry

**Author notes:** Department of Nematology, University of California at Riverside, Riverside, CA, USA. Département de Chimie, Université du Québec à Montréal, Montréal, Québec, Canada (Zoé Joly-Lopez). Department of Plant Biology, Michigan State University, East Lansing, MI, USA. Department of Cell, Developmental, and Regenerative Biology, Icahn School of Medicine at Mount Sinai, New York, NY, USA.

## Abstract

Rice was domesticated around 10,000 years ago and has developed into a staple for half of humanity. The crop evolved and is currently grown in stably wet and intermittently dry agro-ecosystems, but patterns of adaptation to differences in water availability remain poorly understood. While previous field studies have evaluated plant developmental adaptations to water deficit, adaptive variation in functional and hydraulic components, particularly in relation to gene expression, has received less attention. Here, we take an evolutionary systems biology approach to characterize adaptive drought resistance traits across roots and shoots. We find that rice harbors heritable variation in molecular, physiological, and morphological traits that is linked to higher fitness under drought. We identify modules of co-expressed genes that are associated with adaptive drought avoidance and tolerance mechanisms. These expression modules showed evidence of polygenic adaptation in rice subgroups harboring accessions that evolved in drought-prone agro-ecosystems. Fitness-linked expression patterns had predictive value and allowed us to identify the drought-adaptive nature of optimizing photosynthesis and interactions with arbuscular mycorrhizal fungi. Taken together, our study provides an unprecedented, integrative view of rice adaptation to water-limited field conditions.

## INTRODUCTION

The chemical equation for photosynthesis, 6CO_2_ + 6H_2_O + light energy > C_6_H_12_O_6_ + 6O_2_, illustrates that plants cannot maintain high levels of carbon fixation when water availability is limited (Calvin, 1956). In response to environments with restricted or variable water availability, plants have evolved intricate mechanisms to continue to fix resources and maximize survival and seed production (fitness) under drought: 1) drought escape, 2) drought avoidance, and 3) drought tolerance (Levitt, 1980). Escape can be realized through constitutive or drought-induced early flowering, and mechanisms of flowering time regulation have been characterized relatively well (Shrestha et al., 2014). We have recently identified genes, including the transcription factor *OsMADS18*, that contribute to drought escape in rice by studying patterns of covariation between shoot gene expression, flowering time, and fitness (Groen et al., 2020).

However, much less progress has been made in characterizing adaptive variation in drought avoidance and tolerance, because these are complex mechanisms that involve tightly regulated processes at the biochemical, physiological, and whole-plant levels. Drought avoidance may on the one hand involve enhanced water uptake via rapid plastic responses in root hydraulics and/or architecture—a “water-spending” strategy—and on the other hand reduced water loss through changes in leaf area and orientation as well as stomatal conductance—a “water-saving” strategy (Tardieu and Simonneau, 1998). Measurements of stomatal conductance, in conjunction with photosynthetic carbon gain, can be used for determining water use efficiency (WUE; Levitt, 1980). Drought tolerance involves the maintenance of cell turgor through osmotic adjustment or cell wall elasticity, the maintenance of antioxidant capacity, and desiccation tolerance (Levitt, 1980). Plant hormone signaling via abscisic acid (ABA), auxin, and others, plays a vital role in regulating these three drought resistance strategies (Gupta et al., 2020; Todaka et al., 2015).

Rice (*Oryza sativa*) is a staple food for more than 50% of the global population (Wing et al., 2018). Domesticated rice can be subdivided into the *circum*-aus, indica, japonica, and *circum*-basmati subgroups (Huang et al., 2012a; Wang et al., 2018). While the traditional varieties or landraces that make up the temperate japonica, sub-tropical japonica and *circum*-basmati subgroups have predominantly evolved in irrigated agro-ecosystems, landraces in the *circum*-aus, indica and tropical japonica subgroups have also adapted multiple times independently to drought-prone rainfed agro-ecosystems (Gutaker et al., 2020). Finding out the drought resistance strategies that have been selected for in rice, how these strategies are integrated at the whole-plant level, and which regions of the genome regulate them in field settings could inform ongoing efforts to breed and engineer resilient new varieties (Wing et al., 2018).

Traditional forward and reverse genetic mapping approaches, at times combined with RNA sequencing (RNA-seq), have been successful for characterizing the genetic architecture of shoot and root drought resistance traits in carefully controlled plant growth settings, and some candidate genes have been functionally verified (Gupta et al., 2020). Yet, it is not clear how genetic variation in drought resistance strategies relates to the performance of plants at the whole-plant level and how they function in reducing drought damage to fitness-related traits. More insight into how molecular and physiological traits act in concert with morphological traits, and how root and shoot responses to stress are integrated in field environments, is necessary (Des Marais et al., 2012; Groen, 2016; Gupta et al., 2020; Henry et al., 2016). The latter point is of particular importance in light of the growing realization that there is a lab-field gap (Groen and Purugganan, 2016; Poorter et al., 2016; Zaidem et al., 2019). Molecular measurements are typically done in controlled laboratory environments, but plant responses to stress in these circumstances are not fully reflective of how plants react to fluctuating field conditions (Groen et al., 2020; Kawakatsu et al., 2021; Nagano et al., 2012; Plessis et al., 2015; Richards et al., 2012; Swift et al., 2019; Wilkins et al., 2016). Sources of these differences include biotic interactions of plant roots with soil-inhabiting animals and microorganisms such as plant-parasitic nematodes, herbivorous insects, rhizobacteria, arbuscular mycorrhizal (AM) fungi and endophytic fungi (De Vries et al., 2020; Groen and Purugganan, 2016; Poorter et al., 2016; Zaidem et al., 2019). While these interactions are often kept to a minimum in laboratory environments, they can have decisive modulating effects on how drought affects plants (De Vries et al., 2020; Mbodj et al., 2018).

Here, we take an evolutionary systems biology approach, where we characterize morphological and physiological shoot and root trait variation in a rice diversity panel growing in wet or water-limited field conditions and try to better understand how this variation is tied to fitness. We then study the molecular basis of these traits by identifying gene co-expression modules linked to adaptive trait variation, and test if these modules show hallmarks of longer-term selection within domesticated rice. Finally, we assess the predictive value of biological processes enriched in modules relevant to fitness for identifying unmeasured traits that could have adaptive roles and test these predictions in a subsequent crop season.

## RESULTS

### Rice Showed Genetic Variation for Drought Escape and Avoidance Traits

We assembled a core panel of 22 diverse rice accessions from all main subgroups (Supplemental Figure 1, Supplemental Table 1; Huang et al., 2012a; Wang et al., 2018). Most were landraces that have evolved in stably wet irrigated lowland agro-ecosystems, as well as more drought-prone deepwater, rainfed lowland and upland systems, which might be excellent sources of drought resistance-related genetic variants (Groen et al., 2020; Gutaker et al., 2020; Kumar et al., 2014; Torres et al., 2013). Based on previous observations (Groen et al., 2020), we selected accessions for our core panel to have narrower flowering time and biomass windows around the population means than our larger diversity panel with the aim of preventing the strong link between drought escape and plant fitness from overshadowing signals for drought avoidance or tolerance (Supplemental Figure 1).

We then planted identical sister populations (four biological replicate plots per accession) in two lowland fields in the Philippines in the 2017 dry season: a continuously wet field (flooded paddy), and a field where plants were exposed to intermittent drought in the vegetative and reproductive stages (Figures 1A and 1C, Supplemental Figure 2, Supplemental Tables 2 and 3). One accession, Hsinchu 51, displayed dwarfed growth and yellowing, even when not water-limited, and was excluded from all analyses. We studied drought escape by measuring flowering time, and avoidance through measuring a series of morphological and physiological root and shoot traits (Table 1). For root traits, this involved studying the xylem sap exudation rate (Henry et al., 2012), and measuring root density through reconstruction of crown root architecture with the Digital Imaging of Root Traits (DIRT) platform (Bucksch et al., 2014; Das et al., 2015). We also studied drought tolerance by analyzing the biochemical traits of leaf and root osmotic potential (Ψleaf, Ψroot), and the ratio between them (Ψleaf/root). As proxies for plant fitness, we measured the yield of straw and filled grains produced per m^2^ (by dry weight). We also included panicle length and the harvest index (ratio of filled grain dry weight to total shoot dry weight).

**Figure 1.**
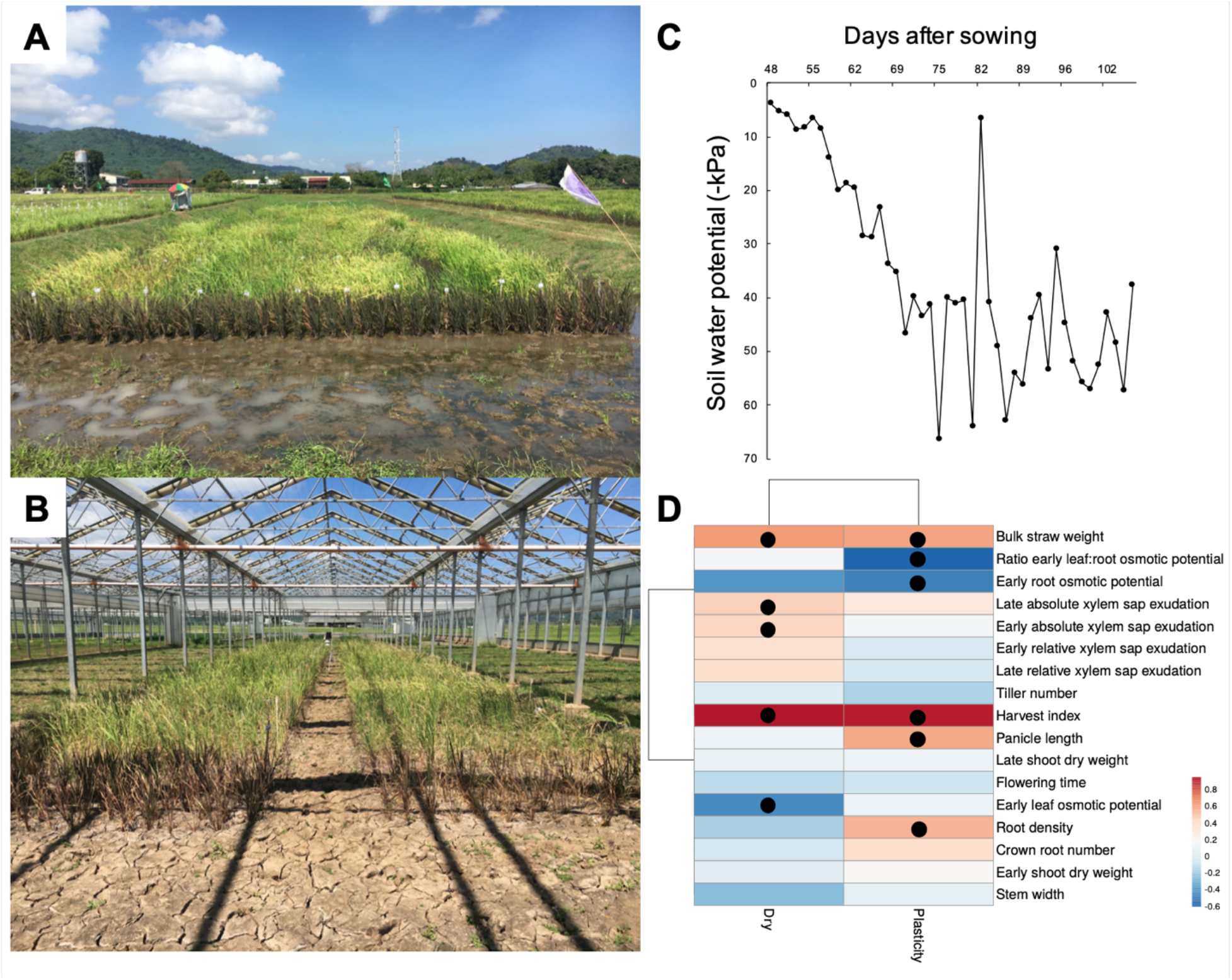
Root physiological parameters are linked to fitness under drought. (A) A panel of 22 diverse rice accessions with representatives of all major sub-populations (planted with four replicates in a randomized complete block design) were monitored for a series of morphological, physiological and molecular traits. (B) The same panel was phenotyped in a rain-out shelter under simulated drought with plots laid out in a design that was identical to the one in the continuously wet paddy. (C) As the season progressed, there was a fluctuating, but increasing, deficit in soil water potential. (D) The heatmap shows Pearson correlation coefficients between baseline and plasticity values of drought resistance traits and the fitness component bulk filled grain weight. Baseline values were assessed for the dry field, and plasticity values by calculating RDPI_s_ between the wet and dry fields. Dots indicate significant correlations (*P* < 0.05).

**Table 1.**
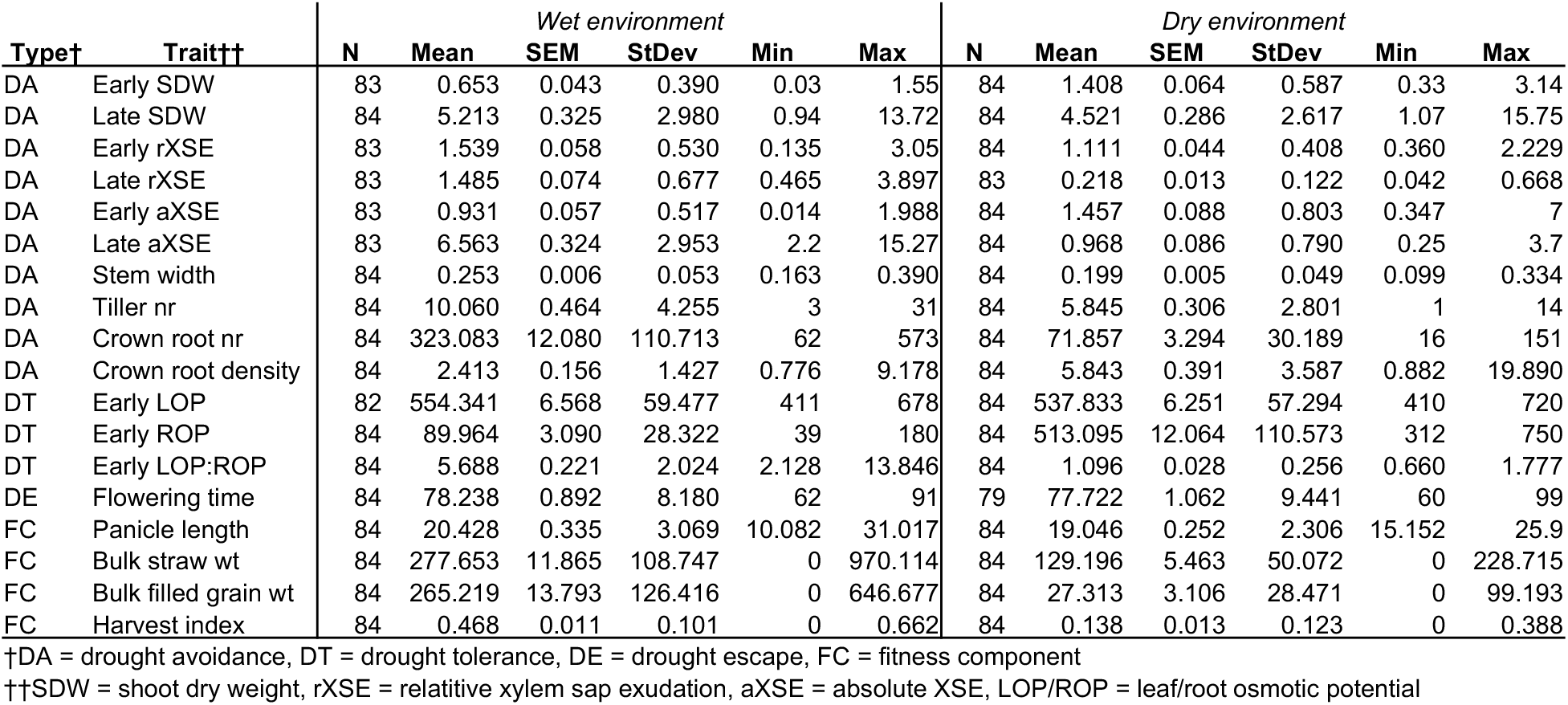
Summary Statistics for Drought Resistance and Fitness Component Traits in the Rice Core Panel across Dry and Wet Environments

| Type† | Trait†† | Wet environment |  |  |  |  |  | Dry environment |  |  |  |  |  |
| --- | --- | --- | --- | --- | --- | --- | --- | --- | --- | --- | --- | --- | --- |
|  |  | N | Mean | SEM | StDev | Min | Max | N | Mean | SEM | StDev | Min | Max |
| DA | Early SDW | 83 | 0.653 | 0.043 | 0.390 | 0.03 | 1.55 | 84 | 1.408 | 0.064 | 0.587 | 0.33 | 3.14 |
| DA | Late SDW | 84 | 5.213 | 0.325 | 2.980 | 0.94 | 13.72 | 84 | 4.521 | 0.286 | 2.617 | 1.07 | 15.75 |
| DA | Early rXSE | 83 | 1.539 | 0.058 | 0.530 | 0.135 | 3.05 | 84 | 1.111 | 0.044 | 0.408 | 0.360 | 2.229 |
| DA | Late rXSE | 83 | 1.485 | 0.074 | 0.677 | 0.465 | 3.897 | 83 | 0.218 | 0.013 | 0.122 | 0.042 | 0.668 |
| DA | Early aXSE | 83 | 0.931 | 0.057 | 0.517 | 0.014 | 1.988 | 84 | 1.457 | 0.088 | 0.803 | 0.347 | 7 |
| DA | Late aXSE | 83 | 6.563 | 0.324 | 2.953 | 2.2 | 15.27 | 84 | 0.968 | 0.086 | 0.790 | 0.25 | 3.7 |
| DA | Stem width | 84 | 0.253 | 0.006 | 0.053 | 0.163 | 0.390 | 84 | 0.199 | 0.005 | 0.049 | 0.099 | 0.334 |
| DA | Tiller nr | 84 | 10.060 | 0.464 | 4.255 | 3 | 31 | 84 | 5.845 | 0.306 | 2.801 | 1 | 14 |
| DA | Crown root nr | 84 | 323.083 | 12.080 | 110.713 | 62 | 573 | 84 | 71.857 | 3.294 | 30.189 | 16 | 151 |
| DA | Crown root density | 84 | 2.413 | 0.156 | 1.427 | 0.776 | 9.178 | 84 | 5.843 | 0.391 | 3.587 | 0.882 | 19.890 |
| DT | Early LOP | 82 | 554.341 | 6.568 | 59.477 | 411 | 678 | 84 | 537.833 | 6.251 | 57.294 | 410 | 720 |
| DT | Early ROP | 84 | 89.964 | 3.090 | 28.322 | 39 | 180 | 84 | 513.095 | 12.064 | 110.573 | 312 | 750 |
| DT | Early LOP:ROP | 84 | 5.688 | 0.221 | 2.024 | 2.128 | 13.846 | 84 | 1.096 | 0.028 | 0.256 | 0.660 | 1.777 |
| DE | Flowering time | 84 | 78.238 | 0.892 | 8.180 | 62 | 91 | 79 | 77.722 | 1.062 | 9.441 | 60 | 99 |
| FC | Panicle length | 84 | 20.428 | 0.335 | 3.069 | 10.082 | 31.017 | 84 | 19.046 | 0.252 | 2.306 | 15.152 | 25.9 |
| FC | Bulk straw wt | 84 | 277.653 | 11.865 | 108.747 | 0 | 970.114 | 84 | 129.196 | 5.463 | 50.072 | 0 | 228.715 |
| FC | Bulk filled grain wt | 84 | 265.219 | 13.793 | 126.416 | 0 | 646.677 | 84 | 27.313 | 3.106 | 28.471 | 0 | 99.193 |
| FC | Harvest index | 84 | 0.468 | 0.011 | 0.101 | 0 | 0.662 | 84 | 0.138 | 0.013 | 0.123 | 0 | 0.388 |

†DA = drought avoidance, DT = drought tolerance, DE = drought escape, FC = fitness component
††SDW = shoot dry weight, rXSE = relative xylem sap exudation, aXSE = absolute XSE, LOP/ROP = leaf/root osmotic potential

**Table 2.**
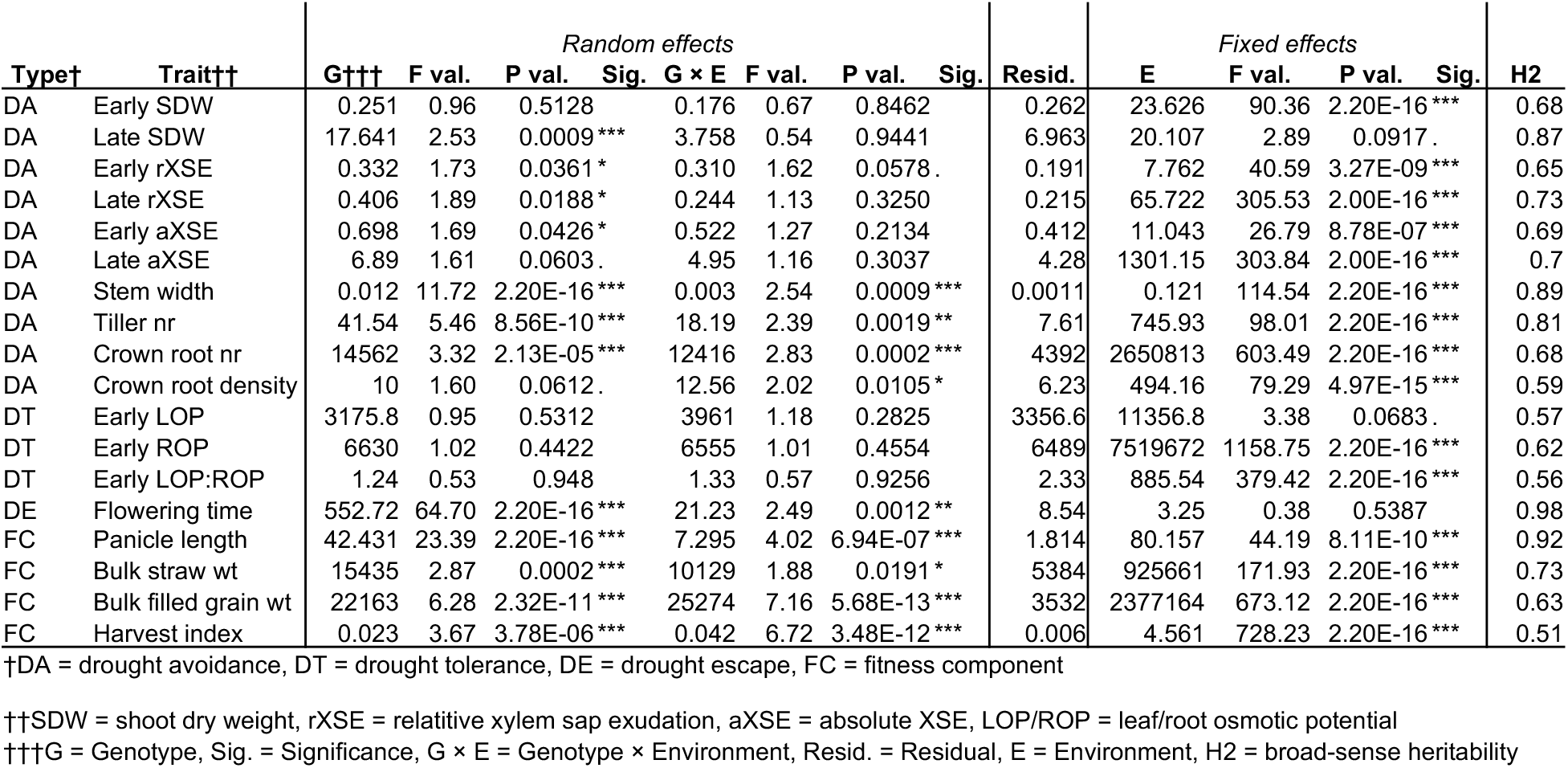
Quantitative Genetic Partitioning of Variation and Significance of Effects across Dry and Wet Environments for Each Trait.

All drought escape and avoidance traits, except early shoot dry weight, generally showed significant genetic variation (*i.e.*, heritability), whereas the drought tolerance traits did not (Tables 1 and 2, Supplemental Table 2). These results show that phenological drought escape traits as well as morphological and physiological drought avoidance traits could be used in breeding if they improve drought resistance. Drought tolerance traits did not show significant genetic variation in our study, presumably owing to higher levels of micro-environmental plasticity in these noisily fluctuating biochemical traits (Henry et al., 2016). However, they could still contribute to drought resistance.

### Several Drought Avoidance and Tolerance Traits Were Associated with Fitness

Next, we tested which traits were genetically associated with measures of plant fitness. Both fitness component traits were significantly positively correlated with one another, as well as with harvest index, in both wet and dry conditions (Pearson’s *r* ≥ 0.6199, *P* ≤ 0.0024). This was also the case for the plasticity of fitness component traits (Pearson’s *r* ≥ 0.5949, *P* ≤ 0.0044). Thus, we focused our analyses for this season on filled grain yield as our fitness proxy (Supplemental Figure 3, Supplemental Table 4).

As expected for this panel chosen to have a narrow flowering time window, drought escape was not significantly linked to fitness: not as flowering time in the dry field (Pearson’s *r* = -0.1303, *P* = 0.5733), and neither as flowering time plasticity (Pearson’s *r* = -0.0955, *P* = 0.6804; Figure 1D). On the other hand, several physiological and morphological traits were indeed linked to fitness under drought. We observed positive fitness correlations for plasticity in panicle length as well as for the drought avoidance traits absolute xylem sap exudation (early and late) and plasticity in root density (Pearson’s *r* ≥ 0.4362, *P* ≤ 0.0481). In particular, high-fitness accessions displayed increased crown root density under drought, despite having lower absolute crown root numbers in these conditions (Table 2). The drought tolerance traits early Ψleaf, and plasticity in early Ψroot and Ψleaf/root showed negative correlations with fitness (Pearson’s *r* ≤ 0.455, *P* ≤ 0.0382; Figure 1D).

### Baseline and Drought-Induced Transcript Expression Show Genetic Variation

We then selected a mini-core of six rice accessions to assess genome-wide gene expression. This mini-core represented the major subgroups of rice that contain accessions from drought-prone deepwater, rainfed lowland and upland agro-ecosystems: tropical japonica (Azucena), indica (Cong Liang 1, IR64 and Kinandang Puti), and *circum*-aus (Bhadoia 303 and Kasalath). The indica accessions come from different sub-populations and agro-ecosystems (Supplemental Table 1). The *circum*-aus accessions respond differently to variation in water availability: Bhadoia 303 shows excellent flooding tolerance as an accession with the SNORKEL1 and -2 haplotypes (Dwivedi et al., 1992; Hattori et al., 2009). Kasalath has the “reference” *SUB1A*, -*B*, and –*C* haplotypes of the submergence-tolerant accession FR13A, and these alleles alter patterns of submergence and drought tolerance (Fukao et al., 2011; Singh et al., 2020; Xu et al., 2006).

We measured transcript levels from leaf blades and crown root tips of four biological replicate plants per genotype per field at 32 days after seedling transplant, and 14 days after withholding water in the dry field, using a liquid automation-based 3’ mRNA-seq quantification approach (Supplemental Table 5; Groen et al., 2020). After filtering out rarely expressed transcripts (expressed in < 10% of shoot or root samples) we included 21,060 shoot transcripts, and 22,707 root transcripts in our analyses (Supplemental Figures 4A and 4B, Supplemental Tables 6 and 7).

**Figure 4.**
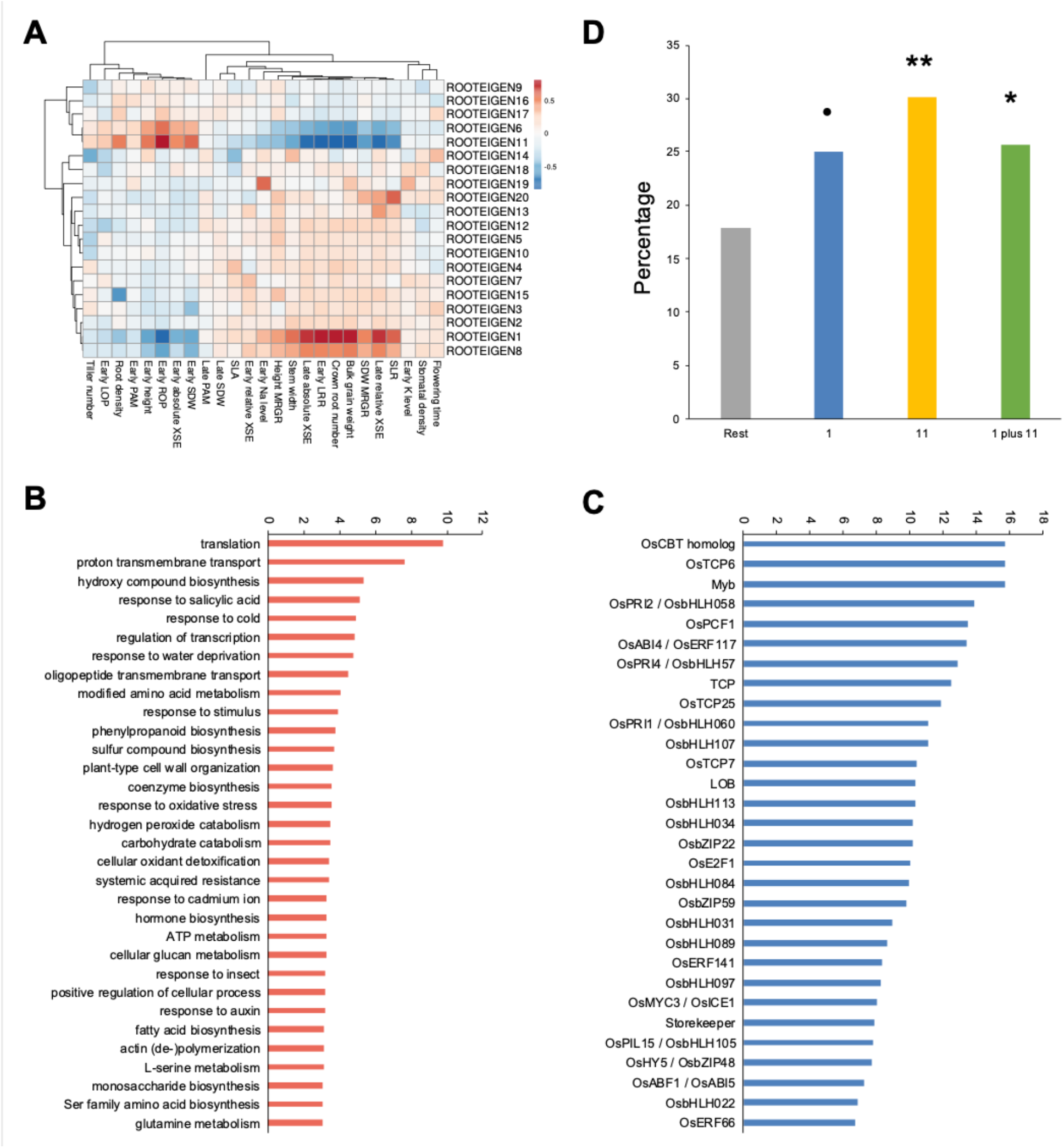
Two fitness-linked root co-expression modules integrate responses to changing abiotic and biotic factors under drought. (A) The heatmap shows Pearson correlation coefficients between the eigengene (PC1) of each of 20 root gene co-expression modules identified through WGCNA and drought resistance traits as well as the fitness component bulk filled grain weight. The 20 modules together represented over 50% of all transcripts included in our analyses. Only modules 1 and 11 were significantly correlated with the fitness component bulk filled grain weight (Bonferroni-adjusted *P* < 2.50×10^-4^). (B) Module 1 was enriched for GO biological processes involved in responses to abiotic and biotic factors (y-axis shows -log_10_ of *P* < 0.05). (C) Promoters of transcripts in module 1 were further enriched for binding sites of several transcription factors (TFs; y-axis shows -log_10_ of FDR *q* < 0.05). (D) Root modules 1 and 11 were enriched for crown root tip transcripts that were differentially regulated in response to interaction with AM fungi in Gutjahr et al. (2015). Notation of significance: ** *P* < 0.01, * *P* < 0.05, • *P* < 0.01.

A principal component analysis (PCA) of all transcripts across all samples revealed a high similarity among the four biological replicates within each genotype × tissue × environment combination. A clear separation between shoot and root samples was observed on the first principal component (PC1), explaining 41% of the total variance (*P* = 6.64×10^-119^; Supplemental Figure 2A, Supplemental Table 8). As expected, the most enriched Gene Ontology (GO) biological process on PC1 was “Photosynthesis” (*P* = 3.00×10^-29^; Supplemental Figure 2B). Field environment was correlated with PC2, explaining 3% of the total variance (*P* = 5.28×10^-14^; Supplemental Table 8) whereas genotype was the third-most important factor, correlating with PC3 (*P* = 4.67×10^-4^; Supplemental Figure 2A).

Separating the data per tissue revealed a mild effect of drought on the shoot transcriptome: we observed 86 differentially expressed genes (DEGs; log_2_ fold change > |1|) across the mini-core accessions. Of these, 44 were upregulated and 42 downregulated after water limitation, while we did not identify unique DEGs for individual accessions (Figure 2A, Supplemental Table 9). Among GO biological processes enriched within shoot DEGs was the pentose-phosphate shunt (Supplemental Table 9), which generates nicotinamide adenine dinucleotide phosphate (NADPH) for reductive synthesis as well as intermediate metabolites for a range of biosynthetic processes (Hou et al., 2007). Pentose-phosphate shunt genes, including *Os6PGDH2* (*OS11G0484500*) are known to be responsive to drought and other abiotic stresses (Hou et al., 2007).

**Figure 2.**
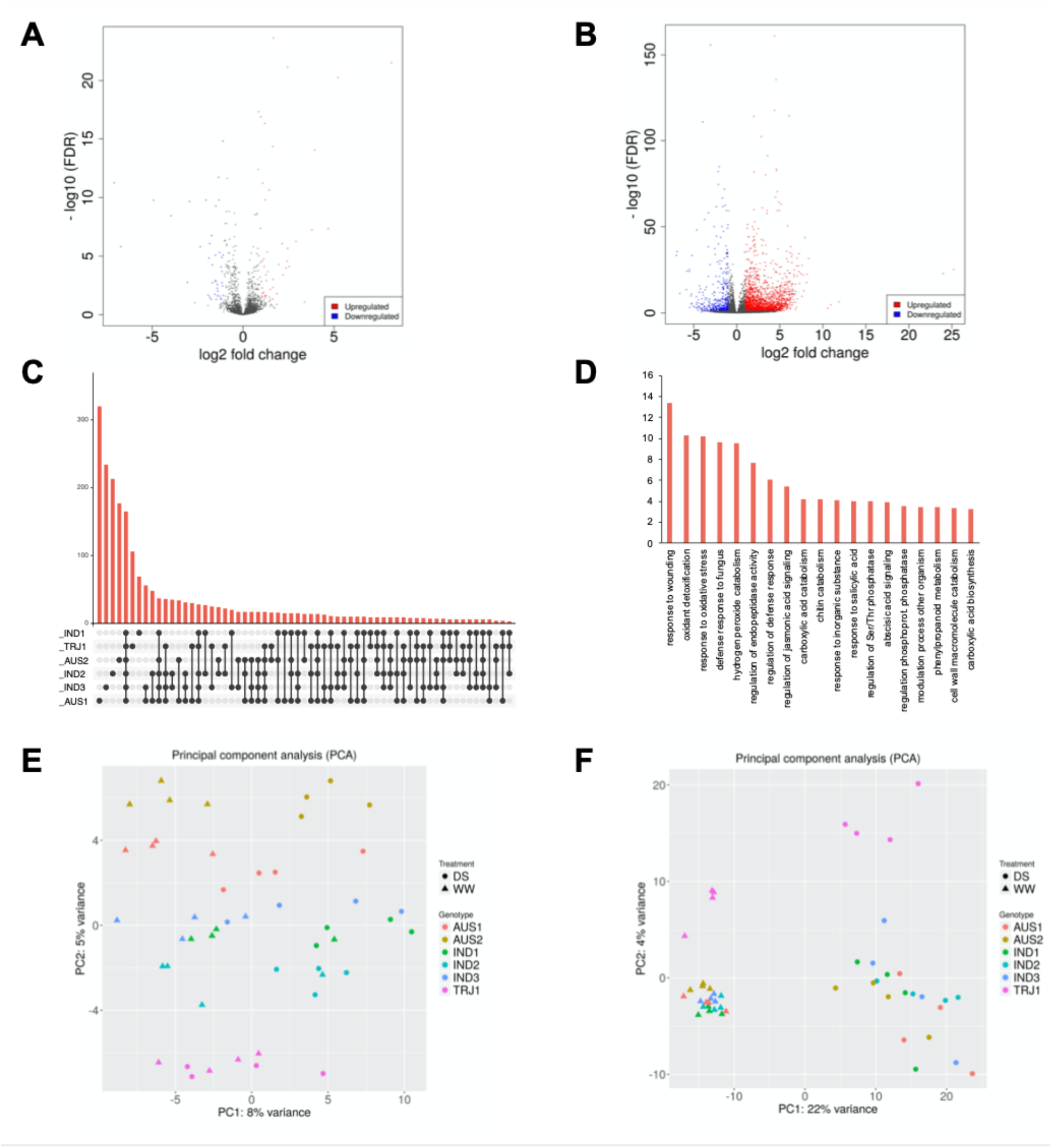
Drought-induced root transcriptional responses were stronger and more accession-specific than shoot responses. (A) The number of DEGs in rice shoot samples (leaf blades) across accessions between wet and dry conditions (FDR *q* < 0.05). (B) Crown root tips under water limitation showed a higher number of DEGs across accessions than leaf blades at the time of sampling (FDR *q* < 0.05). (C) The UpSet plot shows that the largest number of root DEGs (y-axis) was unique for each accession, although a substantial number was shared between all accessions as well. (D) The drought-induced root DEGs shared between all accessions were enriched for GO biological processes related to responses to abiotic and biotic factors (y-axis shows -log_10_ of *P* < 0.05). (E) PCA on the gene expression data shows that PC1 largely separates leaf samples by wet and dry conditions and PC2 largely separates these by accession. (F) Root transcriptome samples separated more clearly by environment along PC1 than shoot samples, while only the roots of the tropical japonica accession Azucena clearly separated from the indica and *circum*-aus accessions along PC2. Abbreviations: WW = wet conditions; DS = dry conditions; AUS1 and -2 = *circum*-aus accessions Bhadoia 303 and Kasalath, respectively; IND1, -2 and -3 = indica accessions Cong Liang 1, IR64 and Kinandang puti, respectively; TRJ1 = tropical japonica accession Azucena.

In contrast to the shoot transcriptome, drought had a much more pronounced effect on the root transcriptome, as reflected by the much higher number of up- and downregulated DEGs (2,158 and 430, respectively) across accessions (Figure 2B, Supplemental Table 10). Individual accessions further varied in the number of drought-responsive DEGs in their roots (Supplemental Figure 4, Supplemental Table 11), with the deepwater *circum*-aus accession Bhadoia 303 from Bangladesh showing the largest number (841 induced and 239 repressed transcripts; Supplemental Figure 4A), and the rainfed lowland indica accession Cong Liang 1 from China the lowest (442 induced and 152 repressed transcripts; Supplemental Figure 4C). These transcriptomic differences might be reflective of known differences in root anatomy and physiology between accessions. For example, lateral root branching in response to drought varies among rice genotypes (Catolos et al., 2017; Kano et al., 2011), and stele and xylem vessel structure as well as sclerenchyma development show genotypic variation (Kondo et al., 2000). Rice varieties further differ in patterns of hydraulic conductivity (Grondin et al., 2016; Henry et al., 2012), as we have measured in our diversity panel (Figure 1D).

To identify which root transcriptional changes constitute a drought response that is conserved across rice sub-populations, we classified root DEGs either as shared between accessions or as unique. We observed that the largest group of shared DEGs was the one consisting of DEGs that all accessions had in common, while much smaller numbers of DEGs were shared between subsets of accessions (Figure 2C, Supplemental Table 11). Among the shared root DEGs we observed an enrichment of biological processes involved in responses to changing water availability such as ones related to ABA signaling and carboxylic acid metabolism (Figure 2D, Supplemental Table 11). Carboxylic acids (especially citric acid) are frequently detected in root exudates and may be able to mobilize phosphorus and other diffusion-limited nutrients that become less available as the soil dries (Gerke, 1995). More unexpectedly, we also found enrichment of processes related to changes in biotic interactions, including interactions with fungi (Figure 2D, Supplemental Table 11). Nutrient uptake under drought may be facilitated further through plant interactions with fungi such as AM fungi (Lanfranco et al., 2018).

We went on to apportion transcript level variance to its sources, so that we could get an impression of the heritability of gene expression patterns. As expected, there was more transcript variation in roots than in shoots, and water availability explained a larger proportion of variance for roots. However, it was surprising to see that genotype explained a similar proportion of transcript variation in both tissues. Among the shoot samples, PC1 explained 8% of the total variance and was correlated with water availability (*P* = 7.96×10^-7^), while PC2 explained 5% of the total variance and was correlated with genotype (*P* = 2.78×10^-34^; Figure 2E, Supplemental Table 8). Among the root samples, PC1 explained 22% of the total variance and was correlated with water availability (*P* = 4.09×10^-27^), while PC2 explained 4% of the total variance and was correlated with genotype (*P* = 3.13×10^-10^; Figure 2F, Supplemental Table 8).

### Shoot and Root Transcript Modules Correlate with Adaptive Drought Resistance Traits

To identify functional gene expression clusters, *i.e.* transcript modules of highly co-expressed genes across plants in the wet and dry environments, we ran weighted gene co-expression network analysis (WGCNA) for the shoots and roots separately (Langfelder and Horvath, 2008). We identified 55 and 112 modules across all shoot and root transcripts, respectively. Of these, 17 shoot and 20 root modules together contained most transcripts (Figures 3 and 4, Supplemental Figure 5, Supplemental Table 12). We focused on these modules for further analysis, and looked for correlations between transcript modules, fitness, and fitness-associated traits.

**Figure 3.**
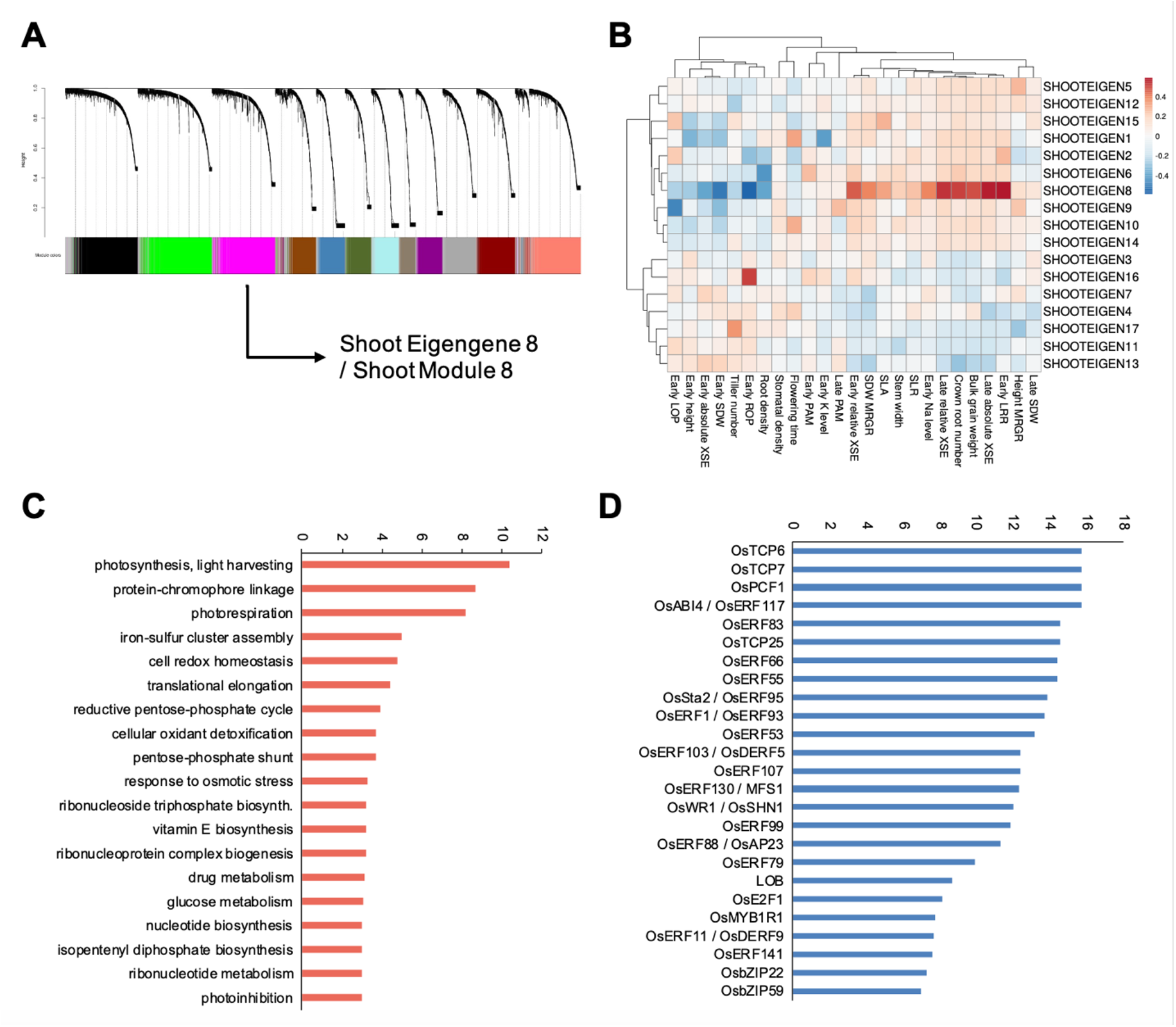
A fitness-linked shoot co-expression module is involved in photosynthesis and regulated by drought-responsive transcription factors. (A) WGCNA identified 17 shoot gene co-expression modules that together represented over 50% of all transcripts included in our analyses. (B) The heatmap shows Pearson correlation coefficients between the eigengene (PC1) of each of the 17 modules and drought resistance traits as well as the fitness component bulk filled grain weight. Only module 8 was significantly correlated with the fitness component bulk filled grain weight (Bonferroni-adjusted *P* < 3.46×10^-4^). (C) Module 8 was enriched for GO biological processes related to photosynthesis (y-axis shows -log_10_ of *P* < 0.05). (D) Promoters of transcripts in module 8 were further enriched for binding sites of several transcription factors (TFs; y-axis shows -log_10_ of FDR *q* < 0.05).

Shoot module 8, as well as root modules 1 and 11, were the only ones that strongly correlated with fitness (filled grain yield) across environments (Bonferroni-adjusted *P* < 3.46×10^-4^ and *P* < 2.50×10^-4^ for shoots and roots, respectively; Supplemental Tables 13 and 14). They also showed significant correlations with all fitness-associated traits: the drought avoidance traits absolute xylem sap exudation (early and late) and root density, and the drought tolerance traits Ψleaf, Ψroot and Ψleaf/root (*P* < 0.05, except for the correlation between shoot module 8 and Ψleaf for which *P* = 0.0852; Figures 3 and 4, Supplemental Tables 13 and 14). Important from evolution and breeding perspectives, all three modules showed robust heritability with *H^2^* ≥ 0.7436 (Supplemental Table 15).

### A Drought-Adaptive Shoot Transcript Module Is Linked to Plant Growth

To gain more insights into the biological roles of the transcript modules associated with drought-adaptive traits, we analyzed whether the modules correlated with additional drought avoidance and tolerance traits that we were able to measure for the mini-core panel. These included the avoidance traits 1) early shoot height, 2) late shoot dry weight, 3) shoot relative growth rates (RGRs, based on height and dry weight), 4) stem-to-leaf ratio, 5) specific leaf area, 6) stomatal density, and 7) chlorophyll fluorescence (early and late). They also included the tolerance traits xylem sap sodium (Na^+^) and potassium (K^+^) content (Supplemental Table 13).

Shoot module 8, as well as root modules 1 and 11, all correlated with the drought tolerance trait xylem sap sodium (Na^+^) content (*P* ≤ 0.0089), and the drought avoidance trait shoot RGR based on dry weight (*P* ≤ 0.0053; Figures 3 and 4, Supplemental Tables 13 and 14). In addition, shoot module 8 and root module 11 correlated with specific leaf area (*P* ≤ 0.0292), while root modules 1 and 11 further correlated with early shoot height (*P* ≤ 1.66×10^-5^), shoot RGR based on height (*P* ≤ 0.0013), and stem-to-leaf-ratio (*P* ≤ 1.82×10^-6^; Figures 3 and 4).

Shoot module 8 correlated with growth-related morphological traits, as reflected in shoot dry weight-based RGR and specific leaf area (Figure 3B), and in addition with water-spending strategy-related physiological traits, as reflected in absolute xylem sap exudation (early and late), Ψroot, and Ψleaf/root (Figure 3B). Growth is in part fueled by photosynthesis and involves increasing both physical size and protein synthesis (Kleessen et al., 2014). Indeed, shoot module 8 was enriched for photosynthesis- and translation-related GO biological processes (Figure 3C, Supplemental Table 16).

Shoot module 8 was further enriched in promoter elements targeted by TEOSINTE BRANCHED1/CYCLOIDEA/PROLIFERATING CELL FACTOR (TCP) and ETHYLENE RESPONSE FACTOR (ERF) transcription factors (Figure 3D, Supplemental Table 16). TCPs are regulators of cell proliferation (Yao et al., 2007), while AP2 ERF TFs such as OsABI4/OsERF117 and OsWR1/OsSHN1/OsERF3 are involved in abiotic stress signaling. Upon drought perception, OsABI4 and OsWR1 transcriptionally activate processes such as protective epicuticular wax production (Mukhopadhyay and Tyagi, 2015; Wang et al., 2012). Other TF binding sites with enrichment included sites for OsbZIP22, OsE2F1, and OsMYB1R1 (Figure 3D). These previously emerged as key regulators of dehydration response- and photosynthesis-related genes in an environmental gene regulatory influence network (Wilkins et al., 2016).

### Two Drought-Adaptive Root Modules Integrate Responses to Abiotic and Biotic Changes

Like shoot module 8, root modules 1 and 11 correlated with growth-related morphological traits as well (Figures 3 and 4). In addition, they were tied to the drought avoidance traits absolute xylem sap exudation (early and late) and root density, which relate to a water-spending strategy. The two root modules were further correlated with the drought tolerance traits Ψleaf, Ψroot, Ψleaf/root and xylem sap sodium (Na^+^) content, which are involved in osmotic adjustment (Chen and Jiang, 2010). These molecular and physiological root traits may be intricately related to phenotypic shoot alterations in response to water limitation. Shoot growth and development depends on a sufficient supply of water coming from the root system, since water spending is of course a prerequisite for photosynthesis (Calvin, 1956). This supply can be promoted by appropriate values of Ψleaf/root and Ψroot (Turner, 2018). Another mechanism known to contribute to water supply to the shoot is root expression of aquaporins (Grondin et al., 2016; Henry et al., 2012; Sakurai et al., 2008). Transcripts from 28 of 34 known aquaporin genes were expressed in our crown root samples (Supplemental Table 17; Nguyen et al., 2013; Sakurai et al., 2005). Root modules 1 and 11, which were not only correlated with plant fitness, but also with xylem sap exudation, were significantly enriched for these aquaporin transcripts (χ^2^ test, *P* = 0.019; Supplemental Table 17). This suggests that root aquaporin expression could contribute to fitness under drought by controlling the water supply to the shoot as measured by xylem sap exudation.

Root module 1 was further enriched for the GO biological processes “response to water deprivation”, “response to hypoxia”, “cellular response to phosphate starvation”, and several metabolism-related processes (Figure 4B, Supplemental Table 17). These processes seem to reflect the differences between the soils in the wet and dry fields and may also be functionally linked to several drought avoidance and tolerance traits to which module 1 was positively correlated: late absolute and relative xylem sap exudation, late crown root number, and Ψleaf/root. In relation to its enrichment for “cellular response to phosphate starvation”, root module 1 was further enriched for “carboxylic acid metabolism” (Figure 4B), a process that may help to mobilize sources of P (Gerke, 1995).

One of the factors that may explain why genes responsive to drought and phosphate starvation make an outsized contribution to between-accession expression differences in root module 1 transcripts is that our mini-core harbors natural genetic variation for possession of the protein kinase PSTOL1 (Chin et al., 2011). For example, PSTOL1 is expressed in the *circum*-aus accession Kasalath, but not in the indica accession IR64 (Chin et al., 2011; Gamuyao et al., 2012). When active, PSTOL1 confers enhanced tolerance to P deficiency, and genes regulated by PSTOL1 activity in *35S::PSTOL1_Kasalath_*-transgenic IR64 plants co-localize with root and drought QTLs (Gamuyao et al., 2012). Strikingly, we observed that root module 1 was indeed enriched for PSTOL1-regulated genes (χ^2^ test, *P* = 0.0135; Supplemental Table 17).

In addition, root module 1 transcripts were regulated more often than expected by chance by the same TCP and OsABI4/OsERF117 TFs that we observed for shoot module 8 (Figure 4C, Supplemental Table 17). There was also a signature of regulation by the TF OsABI5/OsABF1 (Figure 4C), which has known roles in regulating rice responses to water deprivation (Hossain et al., 2010; Zhang et al., 2017; Zou et al., 2008). Compared to shoot module 8 and root module 11, root module 1 was uniquely defined by enrichment for bHLH TFs, including OsPRIs, as well as by enrichment of TF activity from a homolog (*OS03G0191000*) of *OsCBT* (Figure 4C). The latter is a positive regulator of responses to abiotic stress and a negative regulator of biotic interactions (Prasad et al., 2016). OsPRIs regulate root and shoot responses to deficiencies in micronutrients such as iron (Kobayashi et al., 2019; Li et al., 2020).

Root module 11 was enriched for GO biological processes that include “carboxylic acid metabolism”, which was also seen as enriched in root module 1, “response to fungus”, and processes related to oxidative stress responses (Supplemental Figure 4B, Supplemental Table 17). This module was strongly linked with root density (*P* = 4.47×10^-5^), and, compared to shoot module 8 and root module 1, was uniquely enriched for the activity of NO APICAL MERISTEM/ ARABIDOPSIS TRANSCRIPTION ACTIVATION FACTOR/CUP-SHAPED COTYLEDON (NAC) TFs (Supplemental Figure 4C, Supplemental Table 17). One of these TFs was OsNAC2, which is a known regulator of root development, including crown root number (Mao et al., 2020).

Upon closer examination, it was not only root module 11 that was enriched for processes known to be associated with interactions between plant roots and AM fungi. Root module 1 was enriched for such processes as well (Figure 4B; Supplemental Figure 4B), including peptide transport, cell wall organization, and signaling by phytohormones such as auxin and salicylic acid (Gutjahr and Paszkowski, 2009; Gutjahr et al., 2015). Furthermore, root modules 1 and 11 were enriched for transcripts that are differentially expressed in crown roots upon interaction with AM fungi (Gutjahr et al., 2015). Of transcripts in these modules, 25.6% were AM fungus-responsive DEGs compared to 17.9% of transcripts in the rest of the genome (*P* = 0.0427; Figure 4D, Supplemental Table 18). This enrichment was driven in particular by root module 11, of which 30.1% of transcripts were also AM fungus-responsive DEGs (*P* = 0.0014; Figure 4D).

We detected expression of late-stage symbiosis marker genes in 25% of our nodal root samples (Güimil et al., 2005; Gutjahr et al., 2008), with a slight enrichment for plants in dry versus wet conditions (Fisher’s exact test, *P* = 0.0496, Supplemental Table 18). This was despite the fact that we measured gene expression at an early stage after the start of soil drying relative to previously observed progression in the establishment of root interactions with AM fungi, and that we sampled nodal roots, which are typically not colonized as strongly as large lateral roots (Gutjahr et al., 2009; Fiorilli et al., 2015). Among the marker genes were *PT11* (Yang et al., 2012), *STR2* (Gutjahr et al., 2012), *AM25* (Fiorilli et al., 2015), and *AM34* (Fiorilli et al., 2015; Gutjahr et al., 2008), and expression of these was accompanied by relatively frequent expression of common symbiosis marker genes. The latter did not show enriched expression in dry conditions (Fisher’s exact test, *P* = 0.6259, Supplemental Table 18), as expected (Gutjahr et al., 2008). Interestingly, at this relatively early stage we observed marker gene expression in the indica and japonica accessions, but not in the *circum*-aus accessions, which agrees with published patterns of genetic diversity for mycorrhizal symbiosis in rice (Jeong et al., 2015).

Overall, the transcriptome data suggest that the fitness-associated root modules 1 and 11 are not only enriched for transcripts from genes known to be involved in plant responses to drought. They are also enriched for transcripts from genes that mediate root interactions with AM fungi. This brings up the hypotheses that drought may alter plant-fungus interactions and that AM fungi may have an effect on plant fitness in water-limited environments, and we will elaborate further on these below.

### Evidence of Polygenic Selection on Fitness-Associated Root Transcript Modules

Next, we wished to explore if the modules of co-expressed transcripts linked to fitness would show evidence of polygenic selection among rice varieties that regularly experience drought versus rice varieties that do not. One approach was to analyze if genomic regions encoding transcripts within these modules show cumulative effects of differential selection relative to genomic regions encoding the leaf- and root-expressed transcripts outside of these modules. Differential selection may be signified by higher levels of F_ST_, a measure of genetic divergence between groups of individuals (Campbell-Staton et al., 2020; Hämälä et al., 2019). Among rice subgroups, temperate japonica accessions are almost exclusively grown in irrigated agro-ecosystems, whereas *circum*-aus, indica and tropical japonica accessions (including ones in our mini-core) are also grown frequently in drought-prone rainfed systems (Gutaker et al., 2020). We therefore hypothesized that drought-adaptive genome regions should exhibit a tendency to diverge faster between the latter subgroups and temperate japonica than other genome regions.

When we conducted this analysis of polygenic selection we indeed observed above-average levels of F_ST_ between temperate japonica accessions and both the *circum*-aus and the indica accessions in genomic regions encoding the transcripts of fitness-linked co-expression modules compared to genomic regions encoding other transcripts (Welch’s *t*-test, indica: *t* = 1.9944, one-tailed *P* = 0.0231; *circum*-aus: *t* = 2.3723, one-tailed *P* = 0.0089; Figure 5, Supplemental Table 19). Divergence of these genomic regions was not elevated for tropical compared to temperate japonica accessions, although a trend in this direction was visible (Welch’s *t*-test, *t* = 1.3511, one-tailed *P* = 0.0884; Figure 5).

**Figure 5.**
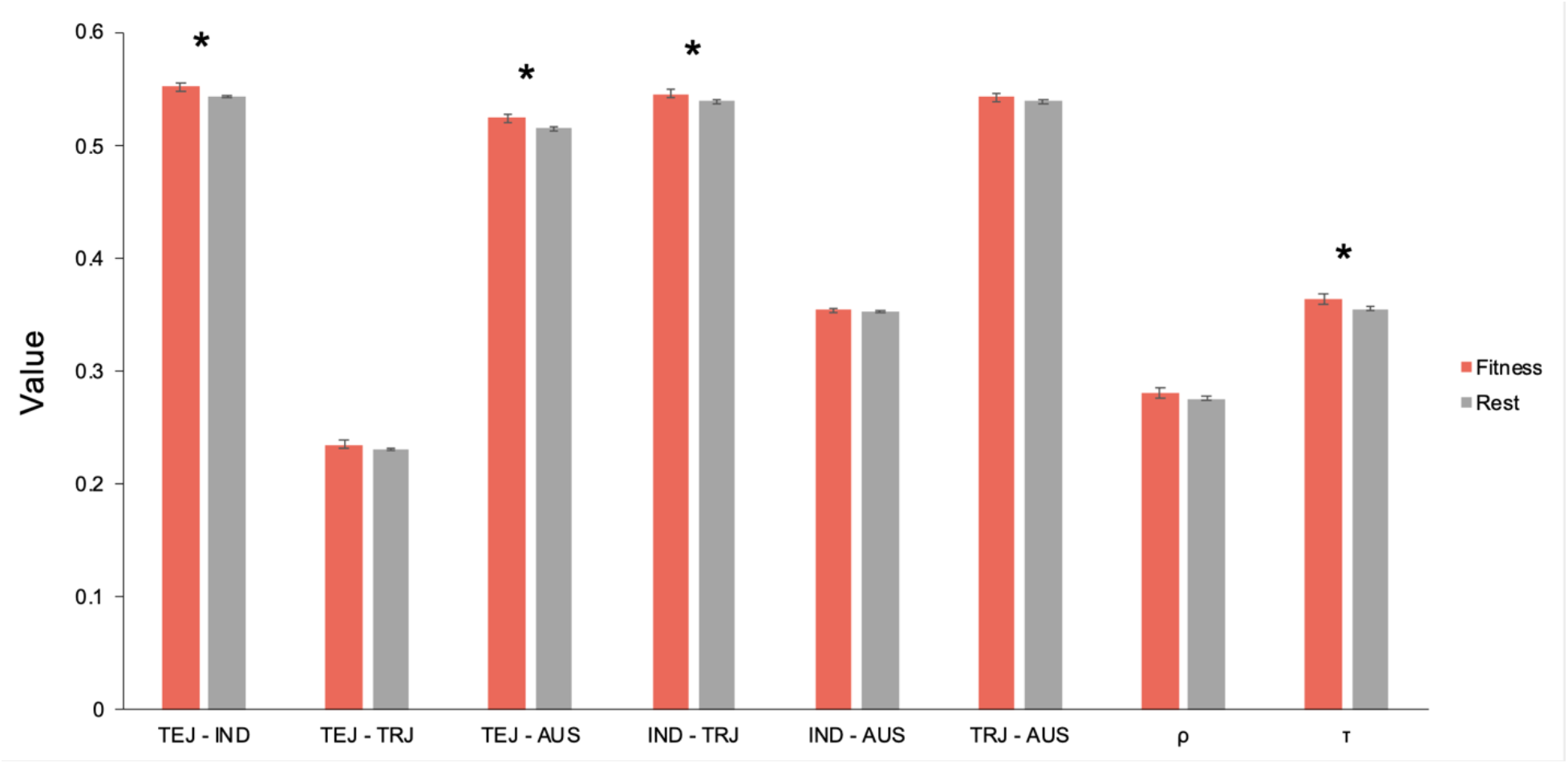
Fitness-linked shoot and root co-expression modules show polygenic signatures of selection across irrigated and rainfed agro-ecosystems. The genomic regions (100-kb windows) encoding the transcripts that are part of the fitness-associated shoot and root gene co-expression modules show higher average values of pairwise population divergence (F_ST_) than other transcriptionally active genomic regions in our experiment in comparisons between temperate japonica accessions (TEJ)—whose members mostly inhabit stably wet irrigated agro-ecosystems—and indica (IND), *circum*-aus (AUS) and tropical japonica (TRJ) accessions, respectively, which are much more regularly found in drought-prone rainfed agro-ecosystems, although the latter difference was not significant. These regions further showed on average higher levels of weak negative or purifying selection (τ), but not of selection in general (ρ), relative to a reference population consisting predominantly of rainfed-environment accessions. Notation of significance: * *P* < 0.05.

We then assessed whether other measures of selection would corroborate our observations on genetic divergence between accessions from drought-prone rainfed compared to stably wet agro-ecosystems. We previously developed GreenINSIGHT, which infers the fraction of nucleotide sites under selection in rice by comparing patterns of intra-species sequence polymorphism with inter-species divergence across dispersed genomic sites, relative to nearby neutrally evolving sites (Gronau et al., 2013; Joly-Lopez et al., 2020). Given the fact that an estimated 89.1% of accessions in GreenINSIGHT’s reference panel were originally collected in drought-prone rainfed agro-ecosystems (Joly-Lopez et al., 2020), we hypothesized that we would observe signatures of selection in the genomic regions from which the transcripts of the fitness-associated co-expression modules originate. We considered two selection-related parameters that GreenINSIGHT computes: ρ (the fraction of sites under any kind of selection) and τ (the fraction of polymorphisms under weak negative selection).

While we observed a trend of higher ρ scores for the genomic regions tied to shoot co-expression module 8 and root co-expression modules 1 and 11 than ρ scores for genomic regions that did not code for transcripts associated with fitness in our experiment, this trend was not significant (Welch’s *t*-test, *t* = 1.1177, one-tailed *P* = 0.1319; Figure 5, Supplemental Table 19). However, we did observe a modestly significant pattern of more pervasive weak negative selection in the form of higher τ scores for these genomic regions (Welch’s *t*-test, *t* = 1.7992, one-tailed *P* = 0.036; Figure 5).

Taken together, these analyses suggest that the genomic regions encoding the transcripts that are part of the fitness-associated shoot and root gene co-expression modules experience polygenic selection in the indica and *circum*-aus sub-populations, whose members more often inhabit drought-prone rainfed agro-ecosystems than the temperate japonica sub-population. Furthermore, weak negative or purifying selection may be more pervasive in these regions compared to other transcriptionally active genomic regions.

### Transcript Modules under Selection Are Predictive for Identifying Additional Adaptive Traits

Patterns of enrichment for biological processes in the shoot and root transcript modules that showed adaptive variation in their behavior gave tantalizing hints of there being unmeasured traits in the 2017 dry season that could have contributed to fitness under drought. For shoot physiological traits, the leaf gene expression data suggested that lower WUE and higher levels of stomatal conductance and photosynthesis might have contributed to higher fitness. In addition, the root gene expression data suggested that such photosynthesis- and fitness-related traits could be linked to root drought avoidance traits. These include xylem sap exudation and environment-responsive developmental plasticity in root density—as deduced from the inferred activity of transcription factors such as OsNAC2 that influence root system architecture. For roots, our field-measured transcriptome data further suggested that there may have been changes in intensity of interactions with AM fungi, which might have come with fitness benefits.

We decided to test if these unmeasured traits from the 2017 dry season could contribute to fitness by measuring them in a second field season in the dry season of 2018 (Figures 6A, Supplemental Table 20). While light- and temperature-related factors were comparable between the two field seasons (Supplemental Figure 6), precipitation- and humidity-related factors showed differences (Figure 6B, Supplemental Table 21). This may explain why not all traits showed significant repeatability (Figure 6C, Supplemental Table 22; Wolak et al., 2012). Repeatability appeared low for biomass-related traits, including the fitness component bulk straw weight (Figure 6C). We therefore decided to consider bulk filled grain weight and bulk straw weight as separate fitness components.

**Figure 6.**
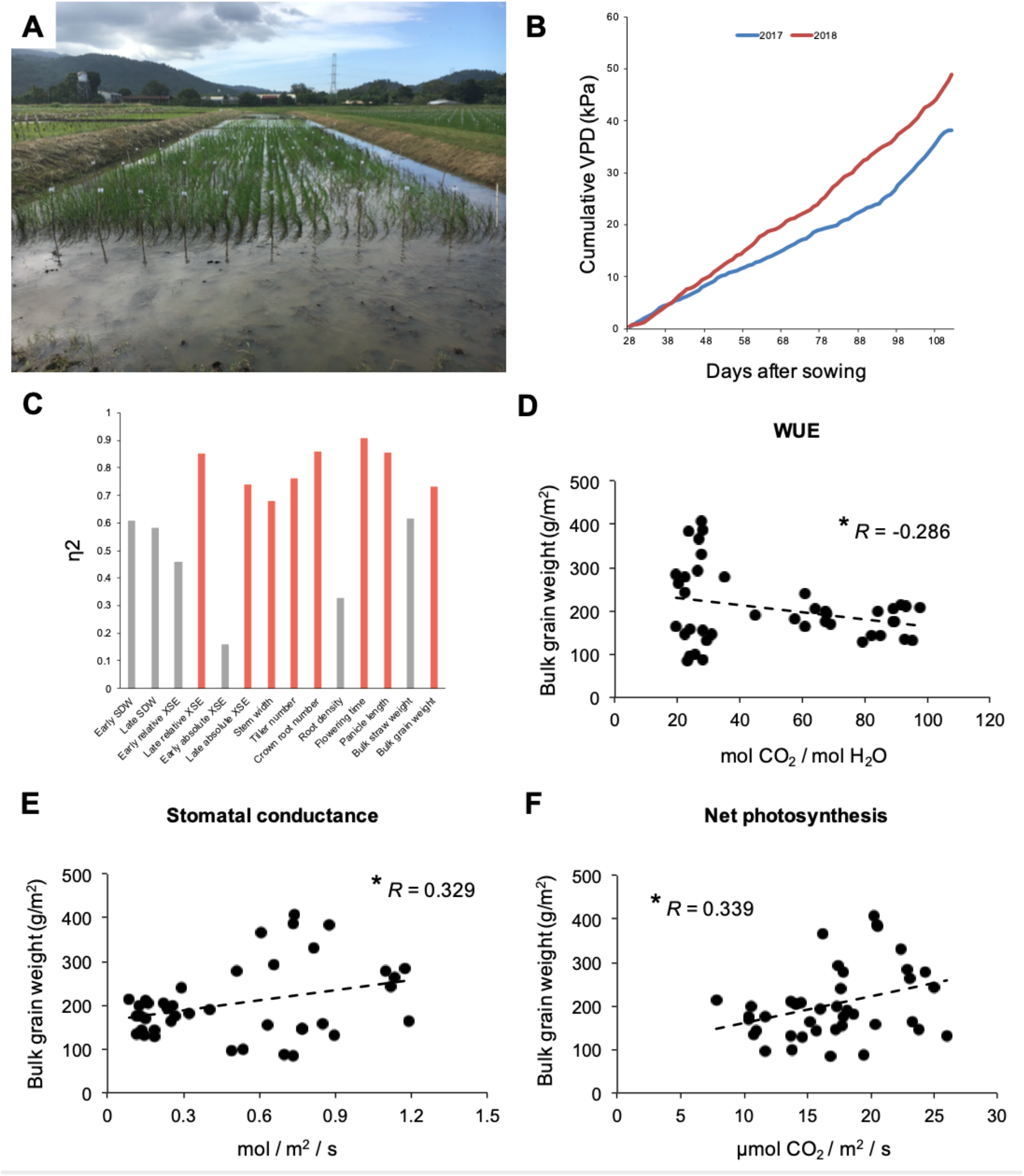
Gene expression patterns predicted links between leaf traits and fitness. (A) The same panel of 22 rice accessions was planted in the same design in wet and intermittently dry field conditions in the 2018 dry season as it was in the 2017 dry season. The incoming clouds symbolize how weather conditions may fluctuate between days and between seasons. (B) Cumulative vapor pressure deficit (VPD) rose more rapidly in 2018 than in 2017. (C) The majority of drought resistance traits showed significant (*P* < 0.05) repeatability (η^2^) between the 2018 and 2017 dry seasons. However, several traits, including some biomass-related traits and root density, did not (gray). (D) WUE showed a significant negative Pearson correlation with the fitness component bulk filled grain weight (*P* = 0.049). (E) Stomatal conductance was significantly positively correlated with grain weight (*P* = 0.0223). (F) Net photosynthesis also showed a significant positive relationship to grain weight (*P* = 0.0184). Notation of significance: * *P* < 0.05.

In shoots, the drought avoidance traits WUE, net photosynthesis and stomatal conductance were linked to bulk grain weight (|*r*| > 0.286, *P* < 0.049; Figures 6D-F, Supplemental Table 23), confirming our expectations based on the gene expression data from the previous dry season. These photosynthesis-related traits are influenced by belowground drought avoidance traits that regulate water transport to the shoots, such as root density and xylem sap exudation, and these root traits showed patterns congruent with our observations for shoots. Drought-induced plasticity in root density was again adaptive in the 2018 dry season (*r* = 0.644, *P* = 0.0016; Figure 7A-E, Supplemental Table 22), despite low repeatability of root density between the 2017 and 2018 seasons (Figure 6C). Furthermore, late xylem sap exudation under drought was correlated with later measurements of net photosynthesis and stomatal conductance in both the 2017 and 2018 dry seasons (Figure 7F), highlighting the links between water transport from the soil and photosynthesis in the leaves.

**Figure 7.**
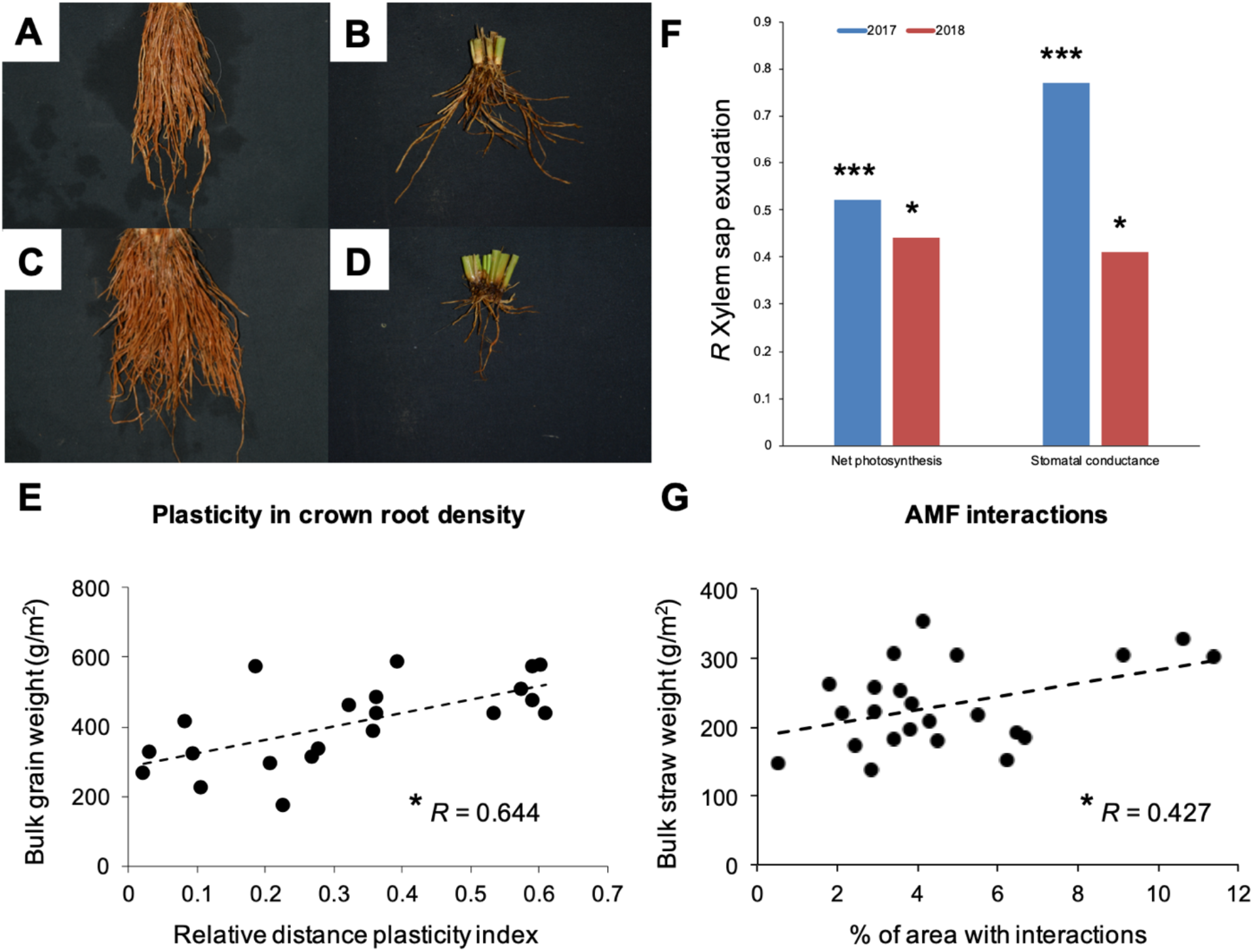
Gene expression patterns predicted links between root traits, leaf traits and fitness. (A) A crown root sample of the indica accession Kinandang puti grown in wet conditions. (B) A crown root sample of the same accession in dry conditions, showing that this accession has limited plasticity in crown root density. (C) The crown roots of the *circum*-aus accession Kasalath showed similar density in wet conditions as those of Kinandang puti. (D) However, the crown root density of Kasalath showed higher levels of plasticity under drought. (E) Plasticity in crown root density was significantly correlated with the fitness component bulk filled grain weight in 2018 (*P* < 0.05), as it was in 2017 (Figure 1). (F) The physiological drought avoidance trait xylem sap exudation showed significant correlations with net photosynthesis and stomatal conductance in the 2017 and 2018 seasons (*P* < 0.05). (G) The intensity of rice root interactions with AM fungi was significantly correlated with the fitness component bulk straw weight (*P* < 0.0374). Notation of significance: *** *P* < 0.001, * *P* < 0.05.

Finally, our expectations for root traits based on predictions from the 2017 field season were confirmed in 2018 as well. Levels of root interactions with AM fungi were tied to bulk straw weight (*r* = 0.427, *P* = 0.0374; Supplemental Table 24), particularly under drought (*r* = 0.853, *P* = 0.0008). AM fungal interactions may further benefit bulk filled grain weight through these effects on biomass as well, although this indirect effect was relatively weak (Estimate = 8.705, *z* = 1.855, one-tailed *P* = 0.032), as estimated through path analysis (Rosseel et al., 2012). Overall, these findings highlighted the power of our gene expression data as a foundation for predicting traits at higher levels of biological organization that may contribute to fitness when rice is faced with limitations in water availability.

## DISCUSSION

The evolutionary systems biology approach we opted to take in this study showed us how plants may maintain fitness under drought through integrating root and shoot physiological responses to adjust hydraulics and photosynthesis. Conducting our measurements of gene expression and physiology in the field allowed us to observe the importance of taking on board changes in biotic interactions when water conditions change to understand plant responses to drought. More field studies of such an integrative nature will need to be done to continue closing the lab-field gap (Groen and Purugganan, 2016; Zaidem et al., 2019).

In our field experiment, we observed fitness associations for a number of traits, including xylem sap exudation and plasticity in root density as drought avoidance traits, as well as Ψleaf and plasticity in Ψroot and Ψleaf/root as drought tolerance traits. Although all tolerance traits showed low heritability, which is a prerequisite for selection to have effects on later generations of plants, the drought avoidance traits showed robust heritability. These findings reinforce previous work showing that xylem sap exudation and root density could make valuable breeding targets, despite the fact that such root-related traits are typically harder to measure in a high-throughput manner than many aboveground traits (Catolos et al., 2017; Henry et al., 2016; Sandhu et al., 2016).

The latter disadvantage might be offset, however, by the fact that we observed stronger heritabilities and trait correlations with gene expression in the root traits compared to the shoot traits. Although many studies have observed the opposite, presumably since root data tends to be noisy and drought stress brings out soil heterogeneity in fields, in our study we may have measured root traits that somehow might be less error-prone than traits that have often been measured in previous studies, such as root length density at depth. Another contributing factor might be the fact that rice shoots tend to stop growing after plants have perceived water limitation during the early vegetative stage and do not show much genetic variation, whereas the roots of the genotypes that perform well under drought are quickly and actively responding to the drought stress, thereby driving a higher degree of genetic variation.

It further seems that rice breeders have had to, and might need to continue to, work against the tendency of natural selection to promote a “water spending” strategy for this species in rainfed lowland conditions (Tardieu and Simonneau, 1998). For example, in our field study with a population that consisted mostly of landraces, xylem sap exudation rate was positively correlated to fitness under drought. However, in several previous studies with populations that contained more breeding lines and different distributions of flowering time and biomass values, the opposite pattern was observed in that drought-tolerant varieties had lower xylem sap exudation rates (Dixit et al., 2015; Henry et al., 2016).

From the literature, it appears that obtaining physiological and, in particular, molecular measurements from the roots of field-grown plants is still limited by considerable practical obstacles compared to obtaining such measurements from shoots. Two published examples provide excellent insights that are extended by our study. In the first, Yu and colleagues measured gene expression in roots of field-grown maize at high and low soil phosphate levels, observing that genes involved in signaling and cell wall metabolism were particularly responsive (Yu et al., 2018). In the second, Kawakatsu and co-workers measured gene expression in both the roots and shoots of a panel of diverse rice accessions grown in relatively dry upland growth conditions (Kawakatsu et al., 2021). The upland agro-ecosystem is a second important rainfed system in addition to the rainfed lowland system we studied (Gutaker et al., 2020; Wing et al., 2018). The authors found that genotypes showed heritable variation in root expression of genes related to auxin signaling and stress responses, and that these processes were linked with root growth characteristics (Kawakatsu et al., 2021).

Our data broadly recapitulates the main findings of these studies, while complementing them in several important ways. Not only do we include an environmental effect, but we also assessed both root and shoot tissues. Crucially, we link measurements of gene expression and physiology with measurement of fitness components, plus analyses of heritability and longer-term effects of selection. That said, in our study, we were not able to include measurements of root growth at depth (*i.e*., beneath a 20-cm depth), which can sometimes explain a large proportion of the drought response in rice (Henry et al., 2011; Uga et al., 2013). Despite this limitation, we found that most of the heritable differences in environmentally sensitive gene expression between accessions could be found in the roots, pointing to the root as the primary response tissue under vegetative stage drought. While the indica accession Cong Liang 1 from China showed the lowest number of DEGs (594 root transcripts), the deepwater *circum*-aus accession Bhadoia 303 from Bangladesh was the most drought-responsive accession with 1,080 root DEGs. Strong plasticity in response to changing water availabilities has been observed for *circum*-aus accessions in previous instances, and it will be particularly interesting to see if these accessions could yield genetic variation that allows for successful plant responses to both drought and flooding (Bin Rahman and Zhang, 2016; Hattori et al., 2009; Fukao et al., 2011; Kano et al., 2011; Sandhu et al., 2016; Xu et al., 2006).

This genetic and environmental variation in gene expression could be summarized in sets of root and shoot co-expression modules. Three of these were significantly tied to fitness and enriched in effects of signaling by phytohormones (including auxin, ABA and salicylic acid) known to mediate plant stress responses (Gupta et al., 2020; Todaka et al., 2015). The modules were further enriched in responses to the nutrient deprivation that comes with water limitation (Swift et al., 2019). Family members of the genes that constitute these modules, such as NAC domain transcription factors and phosphate transporters, appear to be involved in mediating drought-responsive changes in root growth, hydraulic conductivity, and nutrient loading into the xylem across plant species at least as distantly related to rice as the eudicot model plant *Arabidopsis thaliana* (Rosas et al., 2013; Tang et al., 2018).

The two fitness-associated root transcript modules were also enriched for aquaporin-encoding transcripts, which are known to regulate water fluxes towards the shoot under drought stress (Grondin et al., 2016; Henry et al., 2012). In addition, the root modules were correlated with a cluster of co-expressed shoot transcripts enriched in biological processes related to photosynthesis. All three of the root and shoot co-expression modules were intimately linked with the hydraulic traits under selection, which involved xylem sap exudation, root osmotic potential and root density. Taken together, these data suggest positive fitness consequences of coordinated changes in an integrative set of molecular, physiological and developmental traits.

In addition to finding evidence of ongoing selection on the expression of the genes in these modules, we also uncovered signatures of past polygenic selection (Hämälä et al., 2019). The genomic regions underpinning the fitness-associated root and shoot transcript co-expression modules showed elevated genetic divergence (higher levels of F_ST_) between temperate japonica accessions that are typically grown in stably wet, irrigated environments, on the one hand, and indica as well as *circum*-aus accessions that are more often grown in drought-prone rainfed environments, on the other. The genomic regions further appeared subject to more frequent weak negative selection as evidenced by elevated levels of τ, based on a reference population consisting predominantly of rainfed-environment accessions (Gronau et al., 2013; Joly-Lopez et al., 2020). Similar patterns of weak negative or purifying selection have been observed previously in *A. thaliana* for a set of genes that are part of GO biological processes related to biotic interactions (Bakker et al., 2008), for which the fitness-linked co-expression modules we inferred were also enriched, including responses to bacteria, fungi, and insects; responses to chitin and salicylic acid; systemic acquired resistance; and biosynthesis of phenylpropanoids and fatty acids (Supplemental Tables 16 and 17).

Based on these signatures of ongoing and past selection on patterns of gene expression and on variation in physiological and developmental root and shoot traits that transcript levels were tied to, we hypothesized that the transcriptome data would have predictive value for identifying variation in unmeasured traits with consequences for plant fitness. Indeed, a recent study on the rice leaf transcriptome showed that expression changes in response to environmental limitations in water and nutrient availability can have consistent associations with trait variation across seasons (Swift et al., 2019). In keeping with these expectations, our transcriptome data from the 2017 dry season allowed for successful predictions of the importance of photosynthesis-related traits for plant fitness in wet and dry conditions in the subsequent dry season.

Importantly, our transcriptome data extended evidence of the predictive power of gene expression patterns to roots. Drought-induced expression changes that were tied to fitness under drought suggested that plants may respond adaptively to water limitation and concomitant P deficiency by intensifying interactions with AM fungi. Our hypothesis based on these patterns and on previous studies of plant relations with AM fungi (Colard et al., 2011; Lanfranco et al., 2018; Mbodj et al., 2018; Yu et al., 2018), that interactions with AM fungi could have positive fitness consequences under drought, held up in the next dry season. We found that root interactions with AM fungi were positively linked with straw biomass, an important fitness component, within dry as well as between wet and dry field conditions. Future work will be necessary to establish the identities of the fungal interactions partners and to see how these interactions can be harnessed for rice resilience in breeding and agronomy programs (De Vries et al., 2020).

Our evolutionary systems biology approach allowed us to obtain unique insights into how a set of root and shoot traits across levels of biological organization integrates changes in both biotic and abiotic parameters upon drought to benefit rice fitness in lowland field conditions.

## METHODS

### Plant material

A core panel of 22 *O. sativa* accessions was selected from a previously assembled panel landraces and breeding lines (Supplemental Figure 1, Supplemental Table 1). The core included representatives from the *circum*-aus, indica, temperate japonica, (sub-)tropical japonica, and *circum*-basmati subgroups (Groen et al., 2020; Gutaker et al, 2020; Huang et al., 2012a; Wang et al., 2018). Seeds for all accessions were obtained from the International Rice Genebank Collection (IRGC).

### Establishment of field experiments

The first and main field experiment was conducted during the 2017 dry season at the International Rice Research Institute (IRRI) in Los Baños, Laguna, Philippines. Sixteen to twenty-four grams of seed from each of accession was sown onto a seed bed on 1 December 2016, and at 18 days after sowing (DAS) seedlings were pulled and transplanted in a set of two experimental fields. The first one, designated UI (14°008’46.0”N 121°015’51.9”E), remained flooded as a wet paddy. The second field, was located in a rain-out shelter (14°008’33.3”N 121°016’03.4”E) known as UR, remained flooded until irrigation was stopped and the field was drained to start the drought-stress treatment at 36 DAS.

The experiments were arranged in a randomized complete block design with each genotype planted in four replicates with one plant per hill in rectangular, 9 × 5-hill plots with 0.2m × 0.25m spacing for a total of 45 plants per plot. Basal fertilizer was applied at 33 DAS using complete fertilizer (14-14-14) of N_2_, P_2_O_5_ and K_2_O at the rate of 50 kg ha^-1^ each. Manual weeding was conducted regularly in both fields. Kuhol Buster and Actara were applied at 33 and 35 DAS, respectively, to control snail pests, while Actara was applied at 46 DAS and again at 61 and 69 DAS to control insect pests.

To monitor soil moisture levels in the dry field soil water potential was recorded using nine tensiometers (Soilmoisture Equipment) installed at a depth of 30 cm in each replicate. In addition, volumetric soil moisture was recorded at 10-cm depth increments, which was done by frequency domain reflectometry (Diviner 2000, Sentek) through 70-cm PVC tubes installed at nine locations throughout the field.

A second field experiment was set up in the 2018 dry season using the same protocol as much as possible (Supplemental Tables 20 and 21).

### Shoot Trait Measurements

We measured a series of biochemical, physiological, morphological, and phenological traits in shoots to assess differences in drought response between genotypes and individuals. We measured leaf osmotic potential at 49 and 50 DAS (early), and again at 69 and 70 DAS (late) in the wet and dry field, respectively. For this, we collected one leaf per replicate and stored it at −15 °C. The leaves were pressed with a syringe after thawing and 10 μl of sap were pipetted onto a vapor pressure osmometer (Vapro model 5520, Wescor) to measure leaf osmotic potential. We obtained shoot dry weights at 49 DAS (early), and again at 69 DAS (late), in both fields, in the wet and dry field, respectively.

We measured plant height at 49, 55, and 63 DAS as early, intermediate and late time-points, respectively, and calculated the mean RGR using the formula mean RGR = (log*_e_*H*_final_* − log*_e_*H*_initial_*)/*t*, where *t* = time (days) between initial and final measurements of plant height (H; Hunt, 1982). We recorded flowering time as the day on which 50% of plants in a replicate plot flowered.

We selected six accessions as a mini-core for which we obtained additional measurements, including gene expression (see below). For the mini-core accessions we further measured chlorophyll fluorescence on a fully expanded leaf after illuminating at ambient light levels for 20s using the Pulse-Amplitude-Modulation (PAM; MINI-PAM, Walz, Effeltrich, Germany) technique at 50 DAS in both fields (early) and again at 63 and 64 DAS (late) in the wet and dry field, respectively. We calculated the efficiency of Photosystem II or quantum yield (Y_(II)_) as (*F_m_’* – *F_t_*) / *F_m_’*, where *F_m_’* is the maximal fluorescence and *F_t_* the steady-state terminal fluorescence (arbitrary units) (Genty et al., 1989). At 49 DAS (early) we counted the tiller number and measured the leaf area in both fields. For the latter, we used a roller-belt-type leaf area meter (Li-Cor, Model LI-3100C). The leaf area data served as a basis to calculate the specific leaf area (SLA) through dividing by the leaf dry weight.

We further determined the stem-to-leaf ratio for plants in both fields at this time-point, and collected a set of fully expanded leaves for assessing stomatal density. These leaves were first stored in 70% ethanol until we could take epidermal imprints from them at the midpoint of each leaf blade according to Kusumi (2017). For this, we used clear nail polish that we allowed to dry for 10–20 min before removing it with cellophane tape. The imprints were transferred to a microscope slide and imaged at 10× on a BX51 compound microscope fitted with a DP71 camera (Olympus). Images were processed and analysed using ImageJ software version 1.52 (Abramoff et al., 2004), and we counted the number of stomata in areas of ∼0.05 mm^2^ between small veins under a magnification of 200× (Cal et al., 2019; Wang et al., 2016).

We measured additional photosynthesis-related traits for the six mini-core accessions on which we focused our gene expression measurements. For these mini-core accessions we added leaf gas-exchange measurements with a portable LI-6400XT photosynthesis system (Li-Cor Biosciences). We made instantaneous measurements of CO_2_ fixation (*A*) at a photosynthetic photon flux density (PPFD) of 1,000 (2017) or 1,500 (2018) μmol photons m^-2^ s^-1^ and a *C_a_* of 400 μmol CO_2_ mol^-1^, with the flow rate set to maintain a relative humidity of 65%, on one fully expanded leaf per replicate at 86 DAS (late) in the dry field. Before statistical analysis, the leaf gas-exchange data were filtered by excluding data points further than 3SD from the mean.

In the 2018 experiment we measured the same set of biochemical, physiological, morphological, and phenological shoot traits that we measured for all 22 core accessions and the six mini-core accessions in the 2017 experiment. However, in this season we were able to measure the photosynthesis-related traits in both environments, at 79 and 77 DAS (late) in the wet and dry fields, respectively.

### Root Trait Measurements

We further measured a series of biochemical, physiological, and morphological traits in roots to assess differences in drought response between genotypes and individuals. We measured root osmotic potential at 50 DAS (early) in both fields, and again at 69 and 70 DAS (late) in the wet and dry field, respectively. For this, we collected the crown roots of one plant per replicate, blotted them with dry tissue paper, then squeezed out excess water (from the well-watered paddy soil) using a syringe, and stored at −15 °C inside the syringe until frozen. Samples were then thawed, the sap from the root tissue was pressed into a 2-ml tube using the syringe, and the resulting sap was centrifuged at 4,000 RPM for 5 min to remove soil particles before pipetting 10 μl of sap onto a vapor pressure osmometer (Vapro model 5520, Wescor) to measure the root osmotic potential.

We performed measurements of the xylem sap exudation rate (both absolute and relative to shoot dry weight) at 49 and 50 DAS (early), and again at 69 and 70 DAS (late) in the wet and dry field, respectively, according to the protocol described by Morita & Abe (2002) and Henry et al. (2012). Starting at 07:00 h, shoots were cut at a height of around 15cm from the soil surface. Sap emerging from the root zone was collected by covering the cut stems with a 625-cm^2^ cotton towel inside a polyethylene bag that was sealed at the shoot base with a rubber band. After 4h, the previously weighed towel, plastic bag and rubber band were collected and re-weighed immediately to quantify the sap exuded from the intact root system. Shoots were dried and weighed to determine the biomass for each individual for each sampling date. One border row was left between individuals for each sampling date. All xylem sap exudation rate values were both analyzed as absolute measurements and as measurements normalized by the shoot mass of the individual from which sap was collected, in order to account for variation in plant size within and among genotypes.

Concurrent to the xylem sap exudation rate measurements, we sampled the sap exuded from the cut stems of adjacent plants to determine the sodium (Na^+^) and potassium (K^+^) ion content by pipetting sap droplets directly from the cut stems into a 2-mL micro-centrifuge tube. After diluting the samples ∼100-fold to obtain a volume of 10 mL, the samples were analyzed for ion concentration by atomic absorption spectrometry.

We analyzed the architectures of root crowns after excavating the root systems of one plant per replicate at 50 DAS (early) in both fields. Root crowns were excavated using a standard spade to a depth of 25 cm, and a radius of 25 cm around the shoot. We gently washed the root crowns in water before photographing them on a flat, black background accompanied by a scale marker. We then evaluated root traits using the online DIRT platform (Bucksch et al., 2014; Das et al., 2015), focusing on the crown root density.

In the 2018 experiment we measured the same set of biochemical, physiological, and morphological root traits that we measured for all 22 core accessions in the 2017 experiment, plus the additional trait of interaction strength with arbuscular mycorrhizal fungi for the six mini-core accessions on which we focused our gene expression measurements in 2017. We assessed the interaction strength with arbuscular mycorrhizal fungi by performing chitin staining with trypan blue on root crown samples that were taken at 51 DAS (early). Per root system we imaged up to five patches of root area, with each patch ∼0.75 mm^2^ in size, using a BX51 compound microscope fitted with a DP71 camera (Olympus). We inspected these rice root patches for infections by counting the number of intraradical fungal structures with a gridline intersect procedure (Paszkowski et al., 2006; Yang et al., 2012). Before statistical analysis, the AM fungal count data were filtered by excluding data points further than 3SD from the mean.

Trait plasticity was calculated as the as the simplified relative distance plasticity index (RDPI_s_): *P_j_* = |*Z_j,k=2_* – *Z_j,k=1_*| / *Z_j,k=1_*, where *j* is Genotype, *k* is the Focal environment (1 = wet and 2 = dry), *Z* is the Trait value, and *P* is the Trait plasticity (Valladares et al., 2006).

### Fitness or Yield Component Characterization

At physiological maturity, all 7,920 plants that remained in the plots (*i.e.*, individuals that had not undergone destructive sampling during the vegetative stage) were harvested and measured for yield component traits as proxies for plant fitness. These included crown root number as root trait, as well as stem width (cm), tiller number, plant height (cm) and panicle length (cm) as shoot traits. Filled and unfilled grains were sorted and separated from the straw and their bulk dry weights per 1 m^2^ were obtained after drying to 14 % moisture content in an oven at 45°C for three days. The bulked weights of filled grains, unfilled grains and straw were then used to calculate the harvest index as the ratio of filled grain dry weight to the total shoot dry weight (filled and unfilled grain plus straw).

### Tissue Collection for RNA-Seq

Leaf blade and crown root tip sampling was performed in 2017 at 50 DAS (early) on replicate plants over four plots for each of the six mini-core accessions in the wet and dry field from 10:00h to 12:00h (4h after dawn) as described previously (Groen et al., 2020). Four pairs of technicians collected leaf and root tissue in each field, and both fields were sampled simultaneously in the same order by replicate and plot by different teams.

For leaves, two fully expanded leaf blades were selected for sampling. Approximately 12 cm of leaf length was cut into small pieces and submerged into 4 mL chilled RNALater solution in 5-mL screw-cap tubes. For roots, root systems of one plant per replicate in the wet field and four plants per replicate in the dry field were excavated gently using a standard spade. Newly emerging, intact nodal root tips were cut at the base using scissors and submerged into 4 mL chilled RNALater solution in 5-mL screw-cap tubes.

Scissors used for tissue sampling were wiped with 70% ethanol between each plot to avoid contamination. The 96 5-mL tubes with individual tissue samples were placed on ice in styrofoam ice chests, then transferred to a cold room at −4°C overnight. Samples from each of the tubes were then transferred into pairs of 2-mL tubes before being stored at −80°C. One tube of each pair was sent to New York in liquid nitrogen dry shippers to be processed further for mRNA sequencing and long-term storage.

### RNA Extraction, Library Preparation and Sequencing

Frozen leaf blade and root samples were thawed at room temperature and blotted briefly on a KimWipe to remove excess RNALater. The bulk tissue was then flash-frozen in liquid nitrogen and pulverized with pre-cooled mortar and pestle (CoorsTek) to allow extraction of total RNA with the RNeasy Plant Mini Kit according to manufacturer’s protocol (QIAGEN). The RNA was quantified on a Qubit (Invitrogen) before assessing RNA quality on an Agilent BioAnalyzer (Agilent Technologies). The total RNA preps were then stored in nuclease-free water at −80 °C. Total RNA was processed for each individual sample according to a barcoded plate-based 3’ mRNA-seq protocol, as described previously (Groen et al., 2020). This modified version of the SMART-seq2 and SCRB-seq protocols allowed multiplexed pooling of 96 samples before library preparation with the Nextera XT DNA sample prep kit (Illumina). The protocol returned 3’-biased cDNA fragments similar to the Drop-Seq protocol (Macosko et al., 2015). The library consisted of 96 pooled sister samples, *i.e.*, 24 leaf and 24 root samples from the wet field were matched with samples from the same tissues and plot numbers in the dry field. We quantified the cDNA library on an Agilent BioAnalyzer and sequenced it on the Illumina NextSeq 500 in the 2×50-base mode using the following settings: read 1 was 20 bp (bases 1-12 sample barcode, bases 13-20 unique molecular identifier [UMI]), and read 2 (paired end) was 50 bp.

### RNA-Seq Data Processing

The 3’ mRNA-seq reads were quantified according to the Drop-Seq Cookbook from the McCarroll lab using Drop-seq tools v1.12 (J Nemesh, A Wysoker, https://github.com/broadinstitute/Drop-seq/releases) and as described by Groen and co-workers (2020): a wrapper for aligning and parsing not only the reads, but also their embedded barcodes with STAR aligner v020201. STAR used the Nipponbare IRGSP 1.0 (GCF_001433935.1) genome, including plastids, as reference. A reference annotation was generated from Ensembl’s IRGSP nuclear *O. sativa* genome annotation v1.0.37 (ftp://ftp.ensemblgenomes.org/pub/plants/release-37/gff3/oryza_sativa) and supplemented with Refseq’s Mitochondrial and Chloroplast annotations (ftp://ftp.ncbi.nlm.nih.gov/genomes/all/GCF/001/433/935/GCF_001433935.1_IRGSP-1.0). Metadata was generated with Drop-Seq and Picard tools (https://broadinstitute.github.io/picard/). The genome and annotations were indexed using STAR (genomeGenerate with options -- runThreadN 12 --genomeDir inc_plastids --genomeFastaFiles Oryza_sat_CpMt.fa -- sjdbGTFfile 1.0.37_all.gtf --sjdbOverhang 49). Where applicable, annotations were converted between RAP-DB and MSU-7 IDs using the conversion table from the Rice Annotation Project (RAP-MSU_2017-04-14.txt, the latest version can be found at https://rapdb.dna.affrc.go.jp/download/irgsp1.html). For quantification, raw reads were first converted from fastq to unaligned bam format using Picard tools’ FastqToSam and subsequently processed using the unified script (Drop-seq_alignment.sh) in what is essentially default mode for a fastq starting format. Digital gene expression (DGE) profiles were then generated with the DigitalExpression utility using an expected number of barcodes of 96. For QA purposes the DGE profiles were output both as UMI count and raw read count matrices with samples as columns and transcripts as rows. The values represent the number of UMIs that were detected.

To account for differences in the total read number per library, we normalized UMI counts from each sample by dividing by the total number of UMIs detected in that sample. These numbers were multiplied by 1×10^6^ for conversion to transcripts-per-million (TPM). This scaling factor largely represents a consistent in- or decrease across all positive counts in our data matrix. After this, the normalized read counts were subject to blind variance stabilizing transformation provided by the DESeq2 package (Love et al., 2014; R Core Team, 2020).

### Differential Expression

Prior to differential expression analysis the data matrix was split between leaf and root samples. For each organ type samples were analyzed for differentially expressed genes using the DESeq2 package to test for differential transcript expression between each pair of accessions within and among the wet and dry environments using the model: Expression ∼ Genotype + Environment + Genotype × Environment. To detect DEGs, a 5% false discovery rate (FDR) correction for multiple comparisons was determined (Storey and Tibshirani, 2003), and a minimal |1.0| log_2_FC threshold was applied. Contrasts were also generated between the wet and dry environments overall. Despite the lenient threshold for minimum fold change, only few DEGs were detected for leaf-expressed genes between the environments overall, and contrasts between pairs of accessions were only analyzed further for root transcriptomes. For the root samples, private versus common drought-responsive DEGs between accessions were analyzed using Upset diagrams created with the publicly available software Intervene (https://asntech.shinyapps.io/intervene). Principle Component Analyses (PCAs) were performed to explore the gene expression profiles and visualize between-sample distances (Kassambara, 2017).

### Gene Co-Expression Analysis

The variance-stabilized gene expression matrices for leaves and roots were used for the construction of gene co-expression networks using the Weighted Gene Correlation Network Analysis (WGCNA) package (Langfelder and Horvath, 2008), with the soft power parameter (β) set to 6 to ensure that the resulting network exhibited an approximately scale free topology, the P-value ratio threshold for reassigning transcripts between modules set to 0, the cut height of the dendrogram to merge modules set to 0.25, and the minimum size of modules set to 10 transcripts.

### Gene Set Enrichment Analysis

We performed gene set enrichment analysis (GSEA) to gather further biological insight into the DEGs and transcript modules. We considered GO biological processes, using PANTHER’s Overrepresentation Test (released 2021-02-24) with the *O. sativa* genes in the GO database (DOI: 10.5281/zenodo.4495804; released 2021-02-01) as background gene set used to match the foreground set (Mi et al., 2020), as well as TF binding motifs within 300 bp of the transcription start site, using ShinyGO v0.61 (Ge et al., 2020). Enrichment was calculated using Fisher’s exact tests followed by FDR correction. For root co-expression module 11 we added an analysis for enrichment of TF binding motifs to regions within 600 bp of the transcription start site to increase the number of results. For analyses of enrichment in root modules 1 and 11 of genes targeted by PSTOL1 and for genes responsive to interactions with AM fungi we compared our data to sets of genes that were differentially expressed in *35S::PSTOL1Kasalath* transgenic IR64 plants compared to IR64 plants (Gamuyao et al., 2012), and differentially expressed upon crown-root colonization by AM fungi (Gutjahr et al., 2015), respectively. Transcript annotations were converted between RAP-DB and MSU-7 IDs using RAP’s conversion table to make these two datasets compatible with ours (RAP-MSU_2017-04-14.txt, the latest version is available at https://rapdb.dna.affrc.go.jp/download/irgsp1.html).

### Transcript Module Associations with Higher-Level Traits

We identified transcripts associated with the higher-level organismal traits we measured for the populations across the field environments by adding in each higher-level organismal trait in turn to a Pearson correlation matrix of transcript modules using regression models: *Y* = *μ* + *T* + *ε*, where *Y* represents the higher-level trait of interest, *μ* an intercept parameter, *T* denotes the transcript covariate and *ε* residual error. Associations with fitness component traits were deemed significant when they cross Bonferroni-adjusted *P* value thresholds of *P* < 3.46×10^-4^ for shoots and *P* < 2.50×10^-4^ for roots, respectively (Bland and Altman, 1995). Path analysis was performed with the package lavaan version 0.6.6 (Rosseel et al., 2012), which was implemented in R version 4.0.1 (R Core Team, 2020).

### Pairwise Population Divergence Statistics for Transcript Modules

The pairwise population divergence statistics (F_ST_) had been computed as described (Nordborg et al., 2005), using a 100-kb window, between temperate japonica landraces on the one hand, and either *circum*-aus, indica and tropical japonica landraces on the other, respectively (Huang et al., 2010). Values were downloaded from a follow-up study (Huang et al., 2012b).

Variation in F_ST_ across the genome for these sub-populations is of interest in that windows with very high or low values may be seen as candidates for harboring selectively important loci (Akey et al., 2002; Hämälä et al., 2019; Lewontin and Krakauer, 1973), as previously demonstrated for identifying genome regions linked to salinity tolerance in African rice, *Oryza glaberrima* (Meyer et al., 2016).

Each transcript was assigned the F_ST_ value from the genomic region (as a 100-kb window) in which its coding gene was located. We then combined the co-expression module assignments for each transcript with the estimates of sequence evolution to analyze whether selection may be acting differently on the collection of genomic regions giving rise to transcripts that are part of fitness-linked co-expression modules in root and shoot tissue. Estimates of genetic divergence between sub-populations for this collection of genomic regions were acquired by averaging the F_ST_ estimates of the individual regions. These were compared against average F_ST_ estimates from genomic regions harboring root- and/or shoot-expressed genes elsewhere in the genome using Welch’s t-test to account for unequal variances and sample sizes between groups (Welch, 1947).

To find additional evidence of selection on the collection of genomic regions linked to fitness by virtue of giving rise to transcripts that made up shoot module 8 and root modules 1 and 11 we also considered the fraction of sites under any kind of selection ρ and the fraction of polymorphisms under weak negative selection τ of these regions using GreenINSIGHT (Gronau et al., 2013; Joly-Lopez et al., 2020), and comparing these to fractions of sites under selection in other genomic regions using the same approach we used for F_ST_.

### Accession Numbers

Raw RNA sequence data have been deposited as part of SRA BioProject PRJNA564338. Processed RNA expression counts, alongside a key to the RNA sequence data in SRA BioProject PRJNA564338 and the sample metadata, have been deposited in Zenodo under DOI 10.5281/zenodo.4779049. VST-normalized count data can be found in the supplemental material.

## ACKNOWLEDGEMENTS

We thank Leonardo Holongbayan, Eleanor Mico, Nancy Sadiasa, Lesly Satioquia, Bianca Uzziel Principe, and Philip Zamborano for assistance with trait measurements, tissue sampling and field management, IRRI’s Climate Unit staff for providing weather data, the NYU Center for Genomics and Systems Biology GenCore Facility for sequencing support, and NYU High Performance Computing for supplying computational resources. We are grateful to Bruno Guillotin, Ken Birnbaum, Veronica Roman-Reyna, Ricardo Oliva, as well as members of the Purugganan laboratory and IRRI’s Strategic Innovation and Rice Breeding research platforms for insightful discussions. This work was funded in part by grants from the Zegar Family Foundation, the National Science Foundation Plant Genome Research Program and NYU Abu Dhabi Research Institute to M.D.P., a fellowship from the Gordon and Betty Moore Foundation/Life Sciences Research Foundation to S.C.G. (Grant GBMF2550.06), and a fellowship from the Natural Sciences and Engineering Research Council of Canada to Z.J.-L. (Grant PDF-502464-2017).

## Supplemental Figures

**Supplemental Figure 1.**
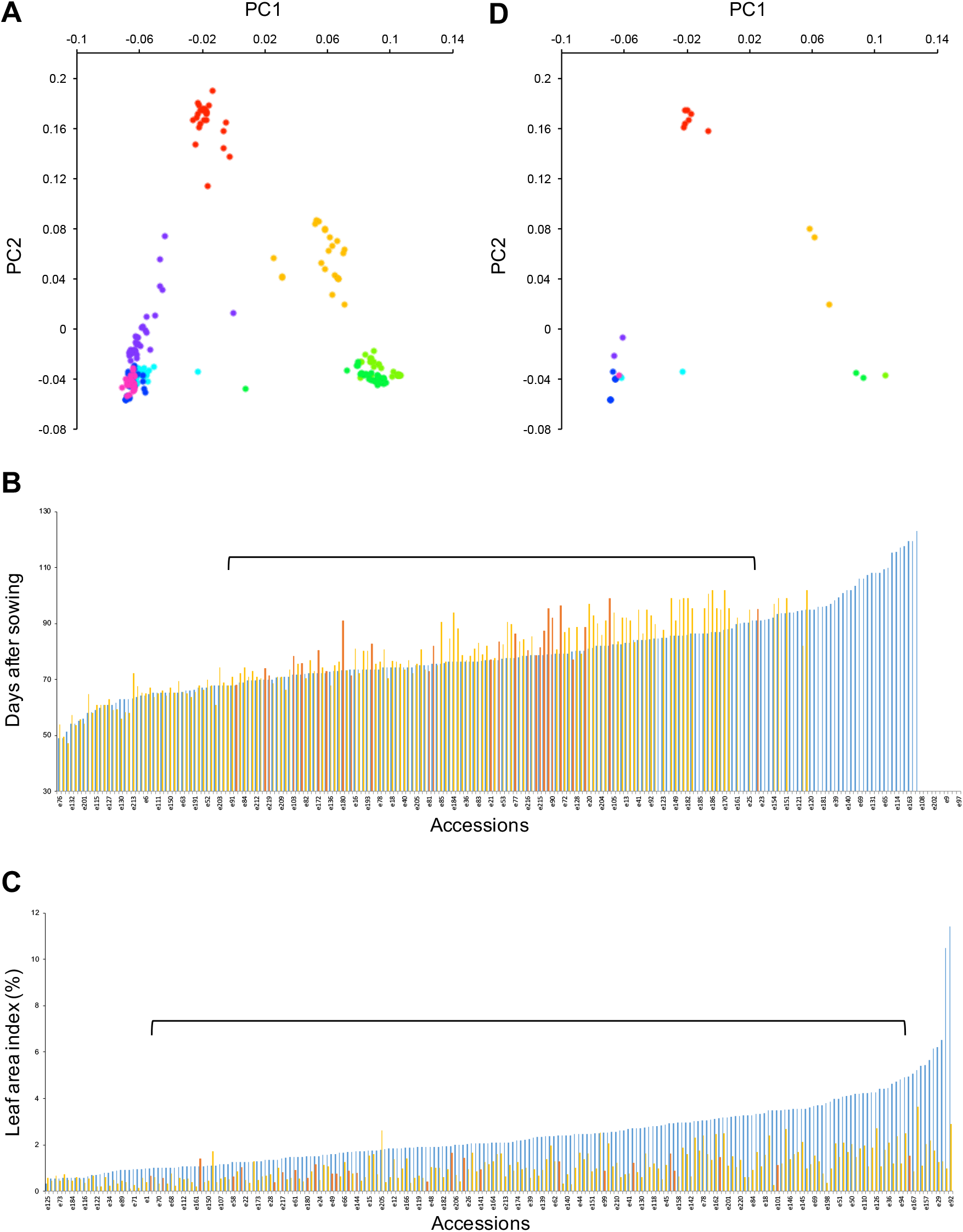
Overview of the core panel of rice accessions selected for this study. (A) PCA for a panel of 215 rice accessions from all major sub-populations that we established previously (Groen et al., 2020). Sub-populations are colored as follows: the *circum*-aus sub-population in red, the *circum*-basmati sub-population in orange, indica sub-populations in blue, magenta and purple colors, and japonica sub-populations in green colors. (B) Distribution of flowering times for the 215 accessions in wet (blue) and dry (yellow) conditions (Groen et al., 2020). Accessions that were selected for our core panel have their flowering times under drought indicated in red underneath the bracket. (C) Distribution of leaf areas for the 215 accessions in wet (blue) and dry (yellow) conditions (Groen et al., 2020). Accessions that were selected for our core panel have their leaf areas under drought indicated in red underneath the bracket. (D) PCA for the core panel of 22 rice accessions selected for planting in the 2017 and 2018 dry seasons. Sub-populations are colored as in (A).

**Supplemental Figure 2.**
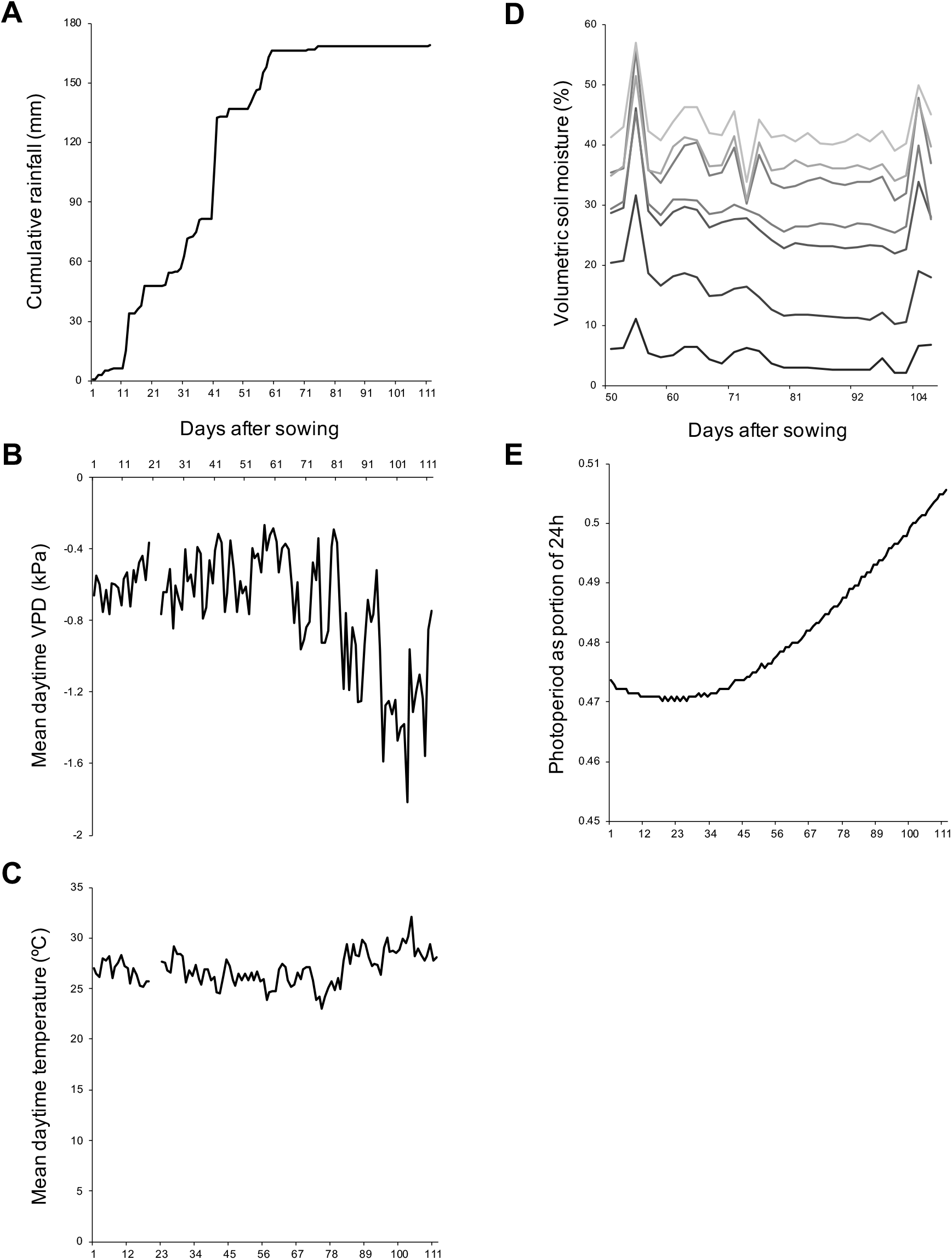
Description of the wet and dry field environments during the 2017 dry season. (A) Cumulative rainfall shows two substantial rainfall events that caused temporary closure of the rainout shelter over the dry field after our rice core panel was transplanted and the drought treatment started (>36 DAS). (B) Patterns of vapor pressure deficit (VPD) suggest that drought became more intense as the season wore on. (C) Patterns of air temperature were consistent with this, as they generally increased over time. (D) Measurements of volumetric soil moisture were in keeping with this as well. Grey lines lighten as data from deeper soil layers are depicted. (E) Days became longer over the course of the season after they had become slightly shorter initially.

**Supplemental Figure 3.**
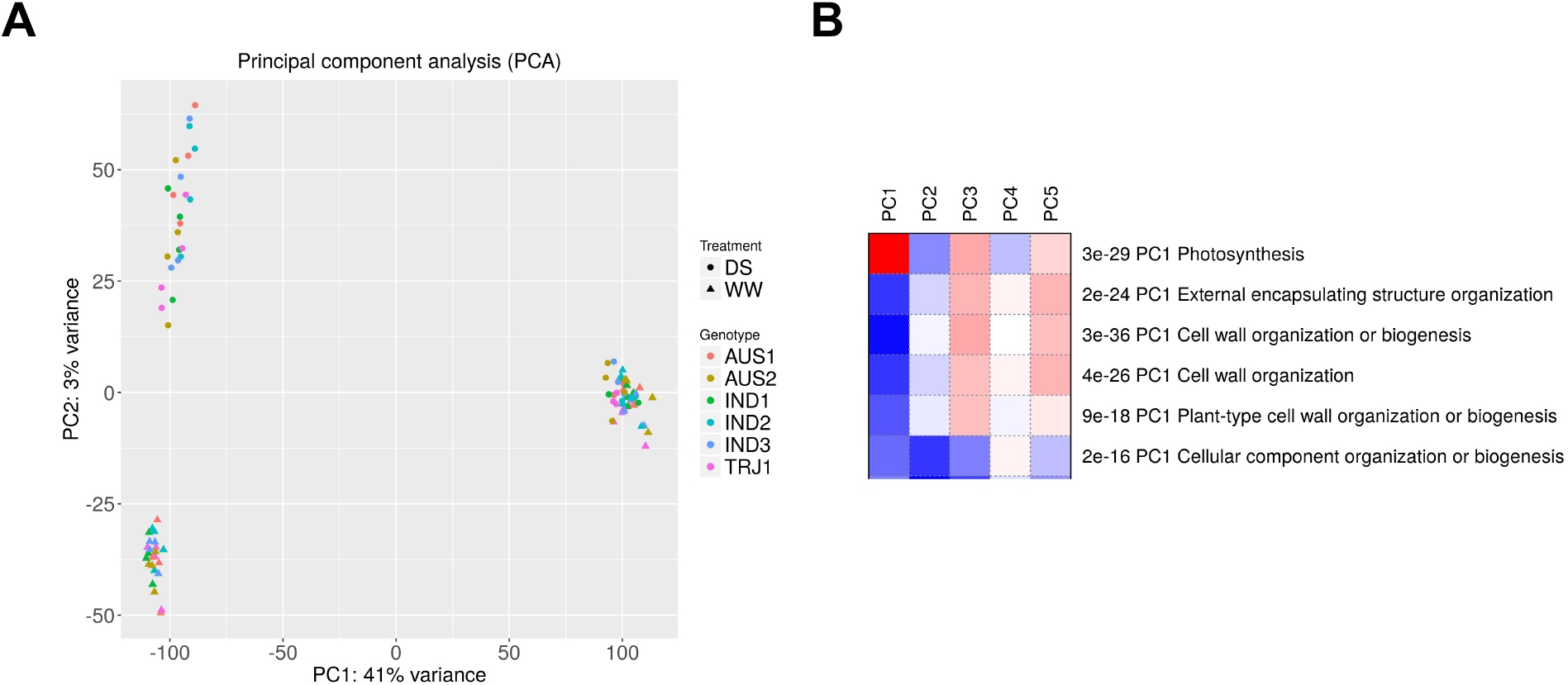
Root and shoot samples of the mini-core panel accessions could be separated based on their genome-wide gene expression profiles. (A) Principal component analysis revealed that shoot samples from leaf blades and root samples from crown root tips were clearly distinct. As expected, they were separated along PC1. Along PC2 we observed a strong distinction between root samples from the wet and dry environments, but not between shoot samples from these environments. (B) Enrichment analysis for gene ontology (GO) biological processes that might drive the distinction between root and shoot gene expression patterns revealed “Photosynthesis” to be the strongest one.

**Supplemental Figure 4.**
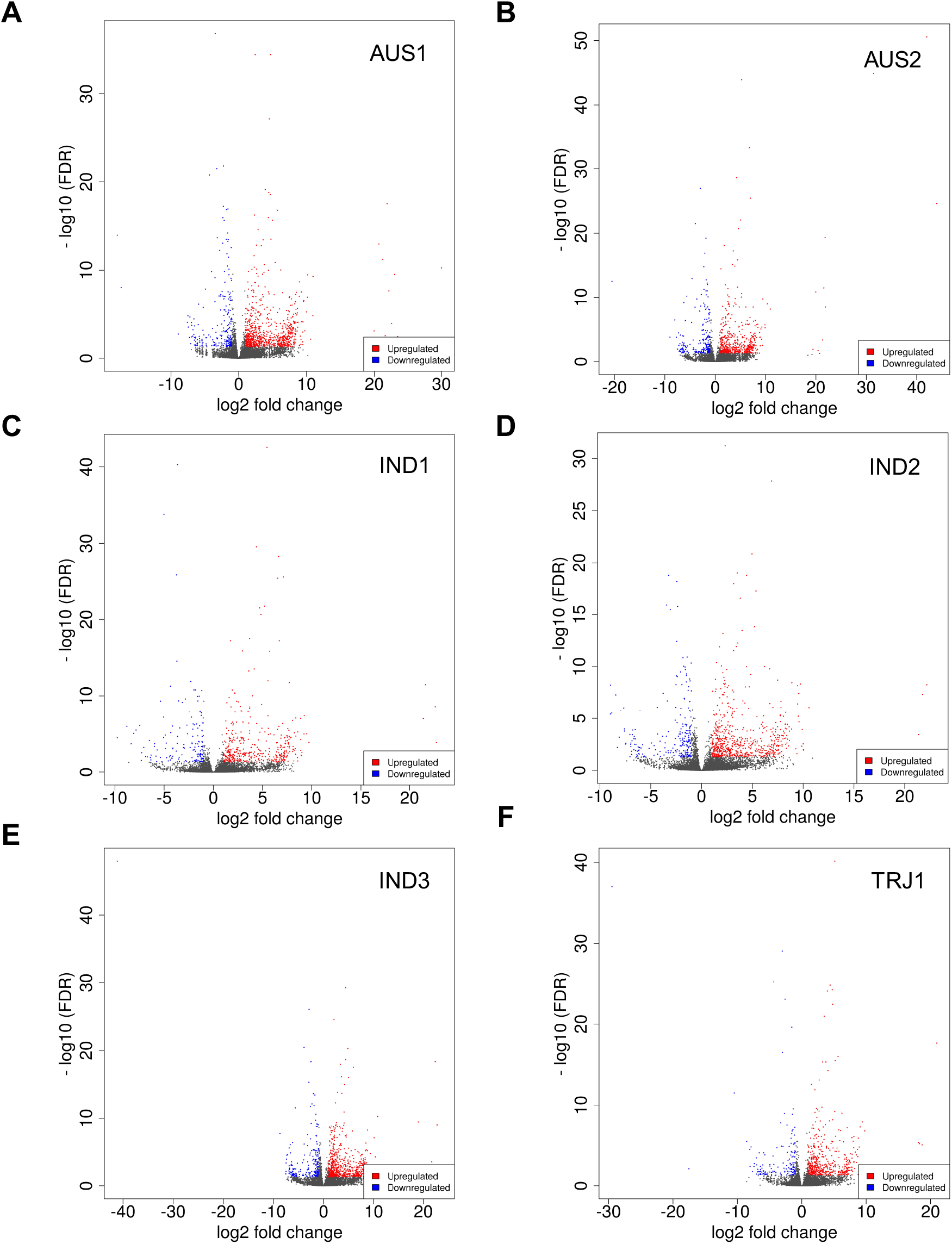
Accession-specific changes to root gene expression in dry versus wet conditions. Crown root tips under water limitation showed varying numbers of drought-responsive DEGs at the time of sampling (FDR *q* < 0.05). (A) The deepwater *circum*-aus accession Bhadoia 303 (AUS1). (B) The rainfed lowland *circum*-aus accession Kasalath (AUS2). (C) The rainfed lowland indica accession Cong Liang 1 (IND1). (D) The irrigated indica accession IR64 (IND2). (E) The irrigated indica accession Kinandang puti (IND3). (F) The upland tropical japonica accession Azucena (TRJ1).

**Supplemental Figure 5.**
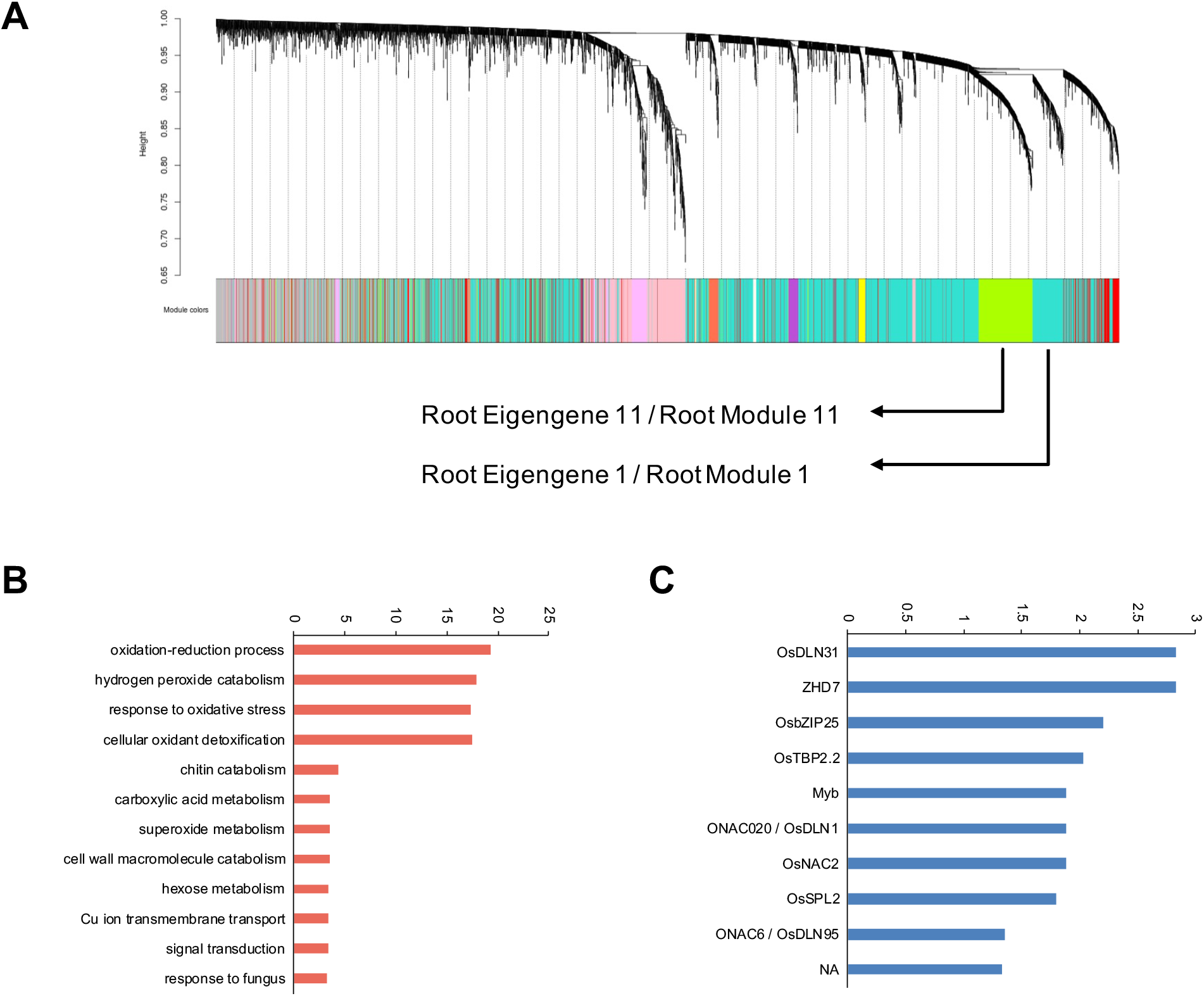
Two fitness-linked root co-expression modules integrate responses to changing abiotic and biotic factors under drought and are regulated by drought-responsive TFs. (A) WGCNA identified 20 root gene co-expression modules that together represented over 50% of all transcripts included in our analyses. Only modules 1 and 11 were significantly correlated with the fitness component bulk filled grain weight (Bonferroni-adjusted *P* < 2.50×10^-4^). (B) Module 11 was enriched for gene ontology (GO) biological process terms involved in responses to abiotic and biotic factors (y-axis shows -log_10_ of *P* < 0.05). (C) Promoters of transcripts in module 11 were further enriched for binding sites of several transcription factors (TFs; y-axis shows -log_10_ of FDR *q* < 0.05).

**Supplemental Figure 6.**
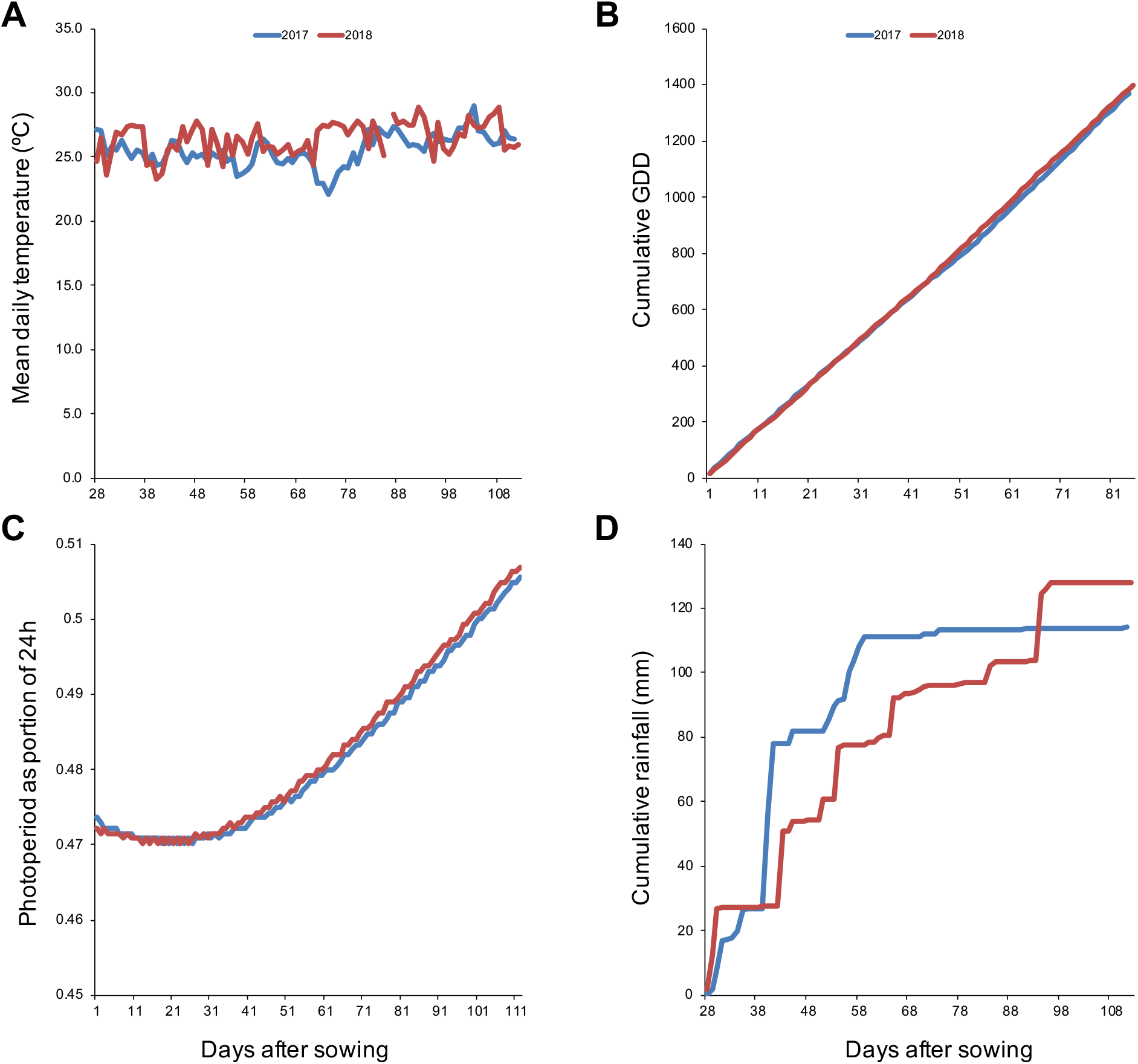
Rainfall and vapor-pressure deficit differed between the 2017 and 2018 dry seasons. (A) Mean daily air temperature showed largely similar patterns between the 2017 and 2018 seasons. (B) The consistent patterns of air temperature between seasons were reflected in the near-identical patterns of growing-degree days (GDDs). (C) Furthermore, and as expected based on the proximity of planting dates, changes in photoperiod progressed similarly between the two dry seasons. (D) On the other hand, cumulative rainfall differed between 2017 and 2018, which matched the patterns of cumulative vapor pressure deficit in Figure 6B.

## Supplemental Tables

**Supplemental Table 1.** List of accessions of the core and mini-core diversity panels.

**Supplemental Table 2.** Experiment timeline, and data on trait as well as fitness component measurements for the 2017 dry season.

**Supplemental Table 3.** Data on weather and soil characteristics during the 2017 dry season.

**Supplemental Table 4.** Correlations between traits, trait plasticity and plant fitness.

**Supplemental Table 5.** Overview of library preparation and transcriptome sequencing for roots and shoots of mini-core accessions.

**Supplemental Table 6.** Normalized transcript level counts for shoots of mini-core accessions.

**Supplemental Table 7.** Normalized transcript level counts for roots of mini-core accessions.

**Supplemental Table 8.** Principal component analyses of RNA-seq samples separated per tissue type and combined.

**Supplemental Table 9.** Differentially expressed genes in rice shoots and enriched GO biological processes among them.

**Supplemental Table 10.** Differentially expressed genes in rice roots.

**Supplemental Table 11.** Overlap between accessions in differentially expressed genes in roots and enriched GO biological processes among overlapping genes.

**Supplemental Table 12.** Root and shoot modules of co-expressed transcripts.

**Supplemental Table 13.** Correlations between traits, fitness and shoot co-expression module eigengenes.

**Supplemental Table 14.** Correlations between traits, fitness and root co-expression module eigengenes.

**Supplemental Table 15.** Effects of genetics and environment on expression variation in transcript co-expression modules.

**Supplemental Table 16.** Gene set enrichment analysis of fitness-linked shoot co-expression module 8.

**Supplemental Table 17.** Gene set enrichment analysis of fitness-linked root co-expression modules 1 and 11.

**Supplemental Table 18.** Expression of marker genes for root interactions with arbuscular mycorrhizal fungi.

**Supplemental Table 19.** Analyses of polygenic selection on fitness-linked root and shoot co-expression modules.

**Supplemental Table 20.** Experiment timeline, and data on trait as well as fitness component measurements for the 2018 dry season.

**Supplemental Table 21.** Comparison of weather characteristics between the 2017 and 2018 dry seasons.

**Supplemental Table 22.** Repeatability of traits between seasons, and correlations between root density and plant fitness components in the 2018 dry season.

**Supplemental Table 23.** Correlations between shoot traits and fitness components in the 2018 dry season.

**Supplemental Table 24.** Correlations between levels of root interactions with arbuscular mycorrhizal fungi and fitness components in the 2018 dry season.

